# Domestic dog lineages reveal genetic drivers of behavioral diversification

**DOI:** 10.1101/2022.04.26.489536

**Authors:** Emily V. Dutrow, James A. Serpell, Elaine A. Ostrander

**Author notes:** Correspondence to: Elaine A. Ostrander, Ph.D., National Human Genome Research Institute, 50 South Drive, Building 50, Room 5351. Bethesda MD, 20892.

## Abstract

Selective breeding of domestic dogs has generated diverse breeds often optimized for performing specialized tasks. Despite the heritability of breed-typical behavioral traits, identification of causal loci has proven challenging due to the complexity of canine population structure. We overcome longstanding difficulties in identifying genetic drivers of canine behavior by developing an innovative framework for understanding relationships between breeds and the behaviors that define them, utilizing genetic data for over 4,000 domestic, semi-feral and wild canids and behavioral survey data for over 46,000 dogs. We identify ten major canine genetic lineages and their behavioral correlates, and show that breed diversification is predominantly driven by non-coding regulatory variation. We determine that lineage-associated genes converge in neurodevelopmental co-expression networks, identifying a sheepdog-associated enrichment for interrelated axon guidance functions. This work presents a scaffold for canine diversification that positions the domestic dog as an unparalleled system for revealing the genetic origins of behavioral diversity.

## Introduction

Selective breeding of domestic dogs to perform specialized tasks may constitute humankind’s most consequential behavioral genetics experiment. Humans have employed dogs for thousands of years: to perform tasks such as herding livestock, killing vermin, hunting, pulling loads, guarding, and companionship (Serpell, 2017; Wilcox and Walkowicz, 1989). To produce dogs that will reliably display traits conducive to executing these functions, humans have selectively bred toward a variety of behavioral ideals. Therefore, genetic analyses of domestic dogs present a unique system for studying how behavioral diversity is biologically encoded, a consequential yet often intractable area of inquiry.

Whereas previous canine genetic mapping and association studies have largely focused on loci underlying behavioral shifts related to domestication or the genetic causes of behavioral issues (Dodman et al., 2016; Duffy et al., 2008; Pendleton et al., 2018; Tang et al., 2014; vonHoldt et al., 2017; Zapata et al., 2016), recent advances in three key domains provide new opportunities to uncover genetic drivers of behavioral traits related to historical working roles (e.g., among gundogs that flush game on command versus terriers that independently track and kill vermin). First, citizen science initiatives related to canine behavior have yielded comprehensive behavioral trait datasets for tens of thousands of domestic dogs (Hsu and Serpell, 2003; Morrill et al., 2022; Salonen et al., 2020; Serpell and Hsu, 2005). Second, the proliferation of publicly available canine whole genome sequencing (WGS) data provided the statistical power necessary to study complex traits, with thousands of genomes representing hundreds of breed types available for interrogation (Bannasch et al., 2020; Jagannathan et al., 2019; Leinonen et al., 2011; Ostrander et al., 2019; Pendleton et al., 2018; Plassais et al., 2019). Third, the development of methods for analyzing ‘Big Data’, both genomic and otherwise, yielded powerful approaches for high-dimensional data exploration that surpass prior capabilities (Amir et al., 2013; Haghverdi et al., 2016; Kuchroo et al., 2022; McInnes et al., 2018; Moon et al., 2019; Street et al., 2018).

Capitalizing on these developments, this study makes three significant advances in the study of canine history and behavior. First, we identify the major canine genetic lineages corresponding to historical breed occupation using a dimensionality reduction approach aimed at capturing and quantifying the local and global complexity inherent in relationships between individual dogs, breeds, and larger canine populations, rather than requiring a hierarchical phylogenetic framework, which is often inconsistent with the nature of dog breed development. Second, we identify behavioral correlates of canine lineage diversification, uncovering key shifts in breed-typical traits conducive to performing specific tasks. Third, we characterize the genetic variants principally driving canine phenotypic specificity, demonstrating the outsized role of non-coding regulatory variation in conferring breed-typical traits and the influence of historical selective breeding for behavioral versus morphological or aesthetic phenotypes.

By studying canine genetic diversification with a lineage-based framework, we circumvent the necessity for tens or hundreds of thousands of genomes to identify complex trait associations across all breeds in aggregate, thereby overcoming the statistical power-related barriers that have stymied recent large-scale efforts to map complex canine behavioral traits (Morrill et al., 2022). As a result, this study uncovers precise neurodevelopmental pathways underlying the major axes of breed diversification and behavioral specification among dogs, revealing how humans shaped the world’s most versatile domesticated animal and establishing canine genetics as a platform for studying the genetic underpinnings of behavioral diversity within a mammalian species.

## Results

### Dimensionality reduction reveals canine lineages corresponding to divergent working roles

To identify genetic drivers of historical occupation-related canine behaviors, i.e., traits conducive to performance of specialized tasks typically assigned to subsets of breeds, we compiled an expansive genomic dataset aimed at maximally representing canine behavioral diversity while accurately capturing global genetic relationships among purebred, mixed breed, free-breeding, and wild canids (Table S1, Table S2). The dataset includes a combination of WGS and single nucleotide polymorphism (SNP) array data from 4,261 individuals, 2,823 of which are purebred dogs representing 226 breeds recognized by the Fédération Cynologique Internationale (FCI; fci.be/nomenclature; Figure 1A, Table S1, Table S2). Owner-reported breed was used as a metric for dataset curation and to aid in visualization; however, the genetic analyses herein are independent of breed membership or recognition. Thus, we included 687 pet dogs of mixed, unknown, or non-recognized breed (Table S1). To capture the full spectrum of selective breeding pressures ranging from purebred dogs to theoretically free-breeding populations, we also included 658 semi-feral village dogs from 47 countries and 93 wild canids from four continents (Table S1).

**Figure 1.**
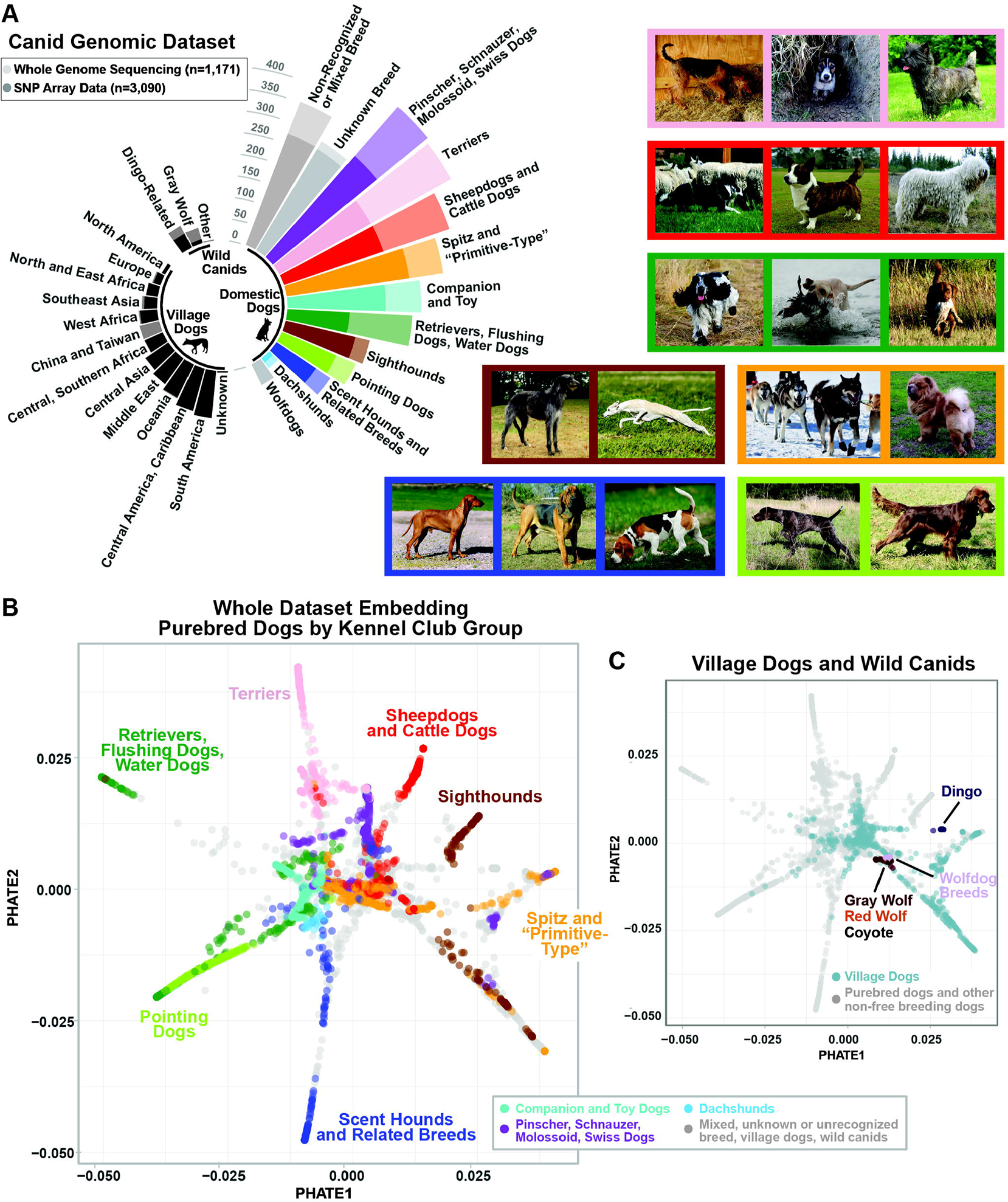
Canid dataset and dimensionality reduction. **(A)** Left: Dataset frequencies by sample and data type grouped by kennel club group designation (Table S2), non-recognized breed, unknown breed, or wolfdog hybrid breeds. Village dogs are grouped by sampling location. Three categories of wild canid are indicated: gray wolves, dingo-related canids, and other (red wolf, coyote). Darker color of each stacked bar indicates number of SNP array samples; upper portion (light color) indicates number of WGS samples. See Table S1. Right: Images of breeds within seven kennel club groups colored according to bar plot. **(B)** PHATE plot of all canid samples colored by group as in panel A. Wild canids and dogs with no kennel club group designation are colored in gray. **(C)** PHATE plot with wild canids and village dogs colored by sample type. All remaining domestic dog samples colored in gray.

We produced a dataset for visualizing and quantifying canine genetic relationships by generating a variance-standardized relationship matrix from a merged dataset of variants from the above samples. While principal component analysis (PCA) alone is often used to visualize genetic data, capturing global trends and outliers, such transformations have a limited ability to capture the sum of all local and global relationships in complex data within two dimensions (Figure S1A). UMAP, a non-linear alternative to PCA, largely clustered individual dogs into isolated groups without preserving global relationships (Figure S1A) (McInnes et al., 2018). To account for global relationships transcending individual breed clusters, we sought to impose no prior assumptions on the structure of our data and instead capture trajectories, linear or bifurcating paths through high-dimensional data that traverse a continuum of states. To do so, we used Potential of Heat Diffusion for Affinity-Based Transition Embedding (PHATE), a tool that encodes local and global data structure in high-dimensional data by transforming relationships among data points into diffusion probabilities that capture the likelihood of transition between states (Moon et al., 2019). We thus generated a maximally informative two-dimensional embedding, i.e., a low-dimensional representation of high-dimensional data for exploratory analysis and visualization. We observed that the PHATE-embedded data formed eight major linear trajectories that emerge from the central portion of the embedding, each roughly corresponding to variance in 2-5 of the input PCs (Figure 1B, Figure S1B). The distal-most portions of the trajectories represent “endpoint” genetic states in the data (i.e., the most divergent genomes within the dataset), with the basal portions of trajectories approximating transitional genetic states between inferred ancestral and endpoint genomes.

We broadly characterized the distribution of purebred dogs in the PHATE embedding using the ten FCI breed group classifications (i.e., kennel club groups broadly categorizing breeds by shared occupation and/or traits; Table S2). We determined that breeds belonging to individual kennel club groups generally separated into distinct trajectories (Figure 1B). One exception was the sighthound group, which split between a largely European-origin sighthound trajectory, and another comprised largely of breeds categorized by FCI as “primitive-type,” which are more aptly described as hunting breeds not originating in Europe or Asia (Figure 1B). FCI groups that did not form one of the major linear trajectories in the two-dimensional visualization formed discrete trajectories observable in three dimensions (Figure S1C). Free-breeding village dogs were largely positioned in the central region of the embedding and wild canids, other than dingo-related populations, clustered tightly near the base of the trajectories comprised of spitz and “primitive-type” breeds, consistent with previous analyses (Figure 1C) (Larson et al., 2012; Parker et al., 2004; vonHoldt et al., 2010). While clusters of breeds previously identified in phylogenetic analyses are generally consistent with the PHATE embedding, PHATE captures the scale of genetic differences between individuals and among related breeds, uncovers ancestral relationships between lineages, accounts for hybridization as a pervasive method of breed creation, and is consistent with multiple major origins for genetic diversity in modern breeds (Figure S1D). For example, the “basal” positioning of the basenji within a single-breed clade paraphyletic to Asian spitz breeds in phylogenetic analyses reflects the basenji’s relative genetic uniqueness among modern purebred dogs, but obscures the genetic relationship the basenji has to other breeds originating in the Middle East and Africa (Larson et al., 2012; Parker et al., 2017; Parker et al., 2004; Plassais et al., 2019; vonHoldt et al., 2010). In contrast, here we preserve complex ancestral relationships, such as that between the basenji, livestock guardian and sighthound breeds originating in the Near East, and village dogs from Africa and the Arabian Peninsula (Figure S1D).

### Canine lineages reflect shifts toward increasingly divergent genetic states

Because annotation based on kennel club-based categories indicated lineage-like trajectories from wild canids to divergent purebred types, we formally characterized the branching patterns observed in the PHATE embedding using pseudotemporal reconstruction, a method for inferring lineages among branching trajectories in high-dimensional data (Street et al., 2018). Pseudotemporal reconstruction applied to canine genetic data serves to order clusters of individuals, in this case partitioned using k-means clustering, into smooth lineages. We designated lineage origin with gray wolves and, in an otherwise unsupervised manner, identified 10 lineages, each of which were named based on breed composition: scent hound, pointer-spaniel, terrier, retriever, herder, sled dog, African and Middle Eastern, Asian spitz, dingo, and sighthound (Figure 2A, S2A; Table S3). We calculated pseudotime, a variable that quantifies positions of data points along lineages without requiring fixed underlying temporal variables such as evolutionary time or a bifurcating branch framework (Street et al., 2018). For example, the herder lineage, as described by pseudotime values for each individual, traverses through livestock-guarding breeds which give rise to the mastiffs along one trajectory and cattle drovers and heelers on another, the latter culminating in sheepdog breeds (Figure 1B, S1D; Figure 2A, S2A). This suggests that sheepdog breeds are the most genetically divergent from the ancestral state of their lineage, and their progenitors may be best approximated by modern European livestock guardian breeds (Talenti et al., 2018). In comparison to divergence time estimations, which are difficult to interpret given the complexity of purebred dog population structure and the nature of breed formation itself, pseudotime values describe each individual in the global context of canine genomes, regardless of the mechanism by which their variation arose.

**Figure 2.**
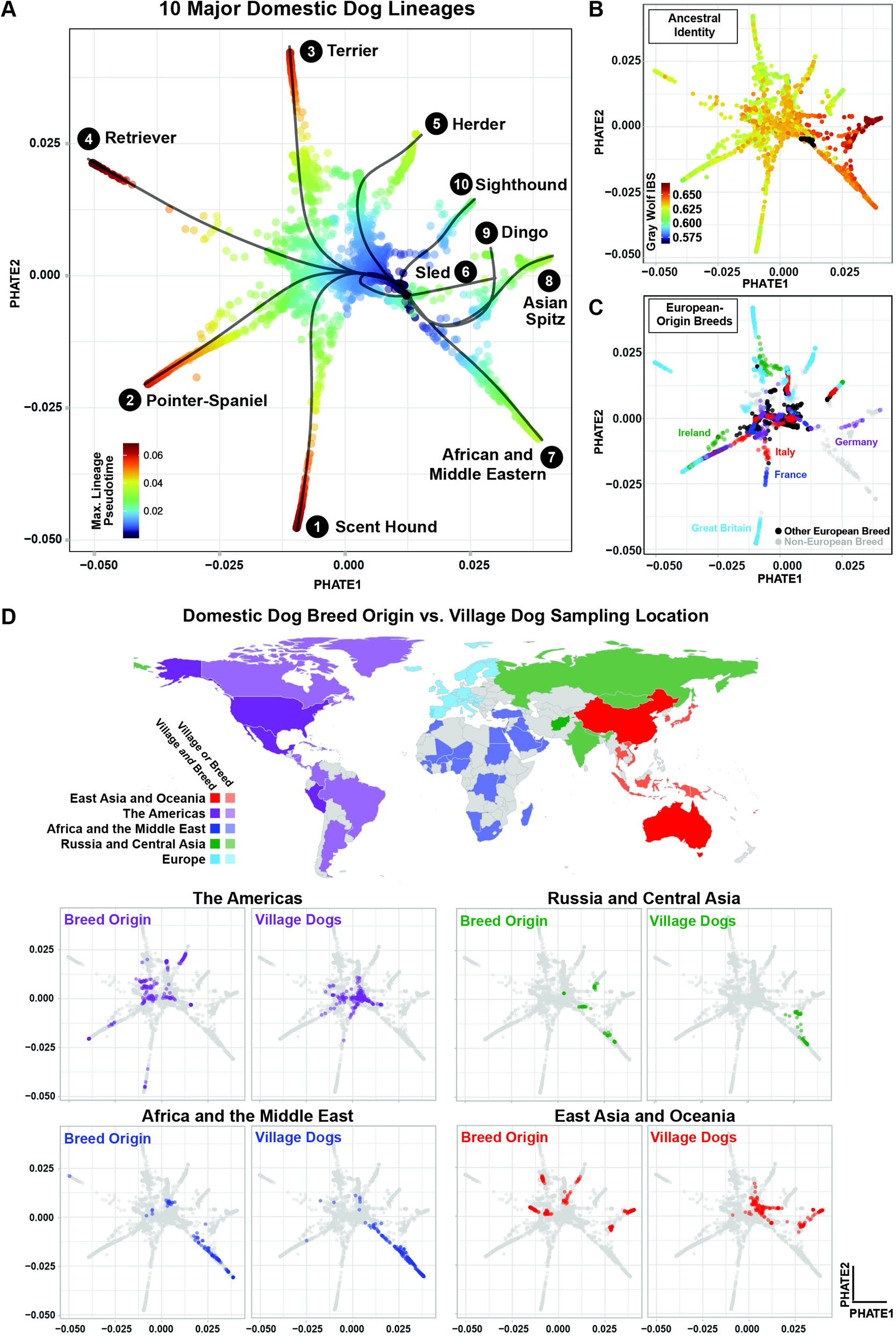
Identification of canine lineages with pseudotemporal reconstruction. **(A)** PHATE plot delineating paths of 10 canine lineages originating with gray wolf. Points are colored by maximum pseudotime in any lineage and plotted lines are principal curves generated with Slingshot (Street et al., 2018). **(B)** PHATE plot showing average pairwise IBS between individuals and each gray wolf. Wolf and coyote samples are colored black. **(C)**. PHATE plot indicating samples from breeds with European origins (non-European origin breeds in gray). **(D)** Top: Geographic origin of breeds (historical, FCI-based) and village dog sampling location. Darker shade indicates locations with breed origin and village dog representation in the dataset. Lighter shade indicates countries for which dataset includes either domestic or village dogs. Bottom: PHATE plots comparing breed origin (left in each pair) to village dogs of the same geographic location (right in each pair) for non-European geographic regions.

For each individual in the dataset, we calculated identity-by-state (IBS) with each gray wolf, observing that lineage structure did not appear to be singularly driven by decreased mean ancestral identity (Figure 2B). Instead, lineage progression was broadly characterized by decreased inter-breed IBS, with terminal breeds (those with higher pseudotemporal positions) displaying the most unique constellation of genotypes compared to all other dogs. For instance, when examining a subset of scent hound lineage breeds, we found that breeds with the highest pseudotemporal positions (e.g., otterhound, bloodhound), had the least amount of shared identity with all other dogs, while breeds with lower pseudotemporal positions generally showed higher shared identity with all other dogs both within and outside the scent hound lineage (Figure S2B).

We next characterized the degree to which genetic relationships among and across lineages were related to geographic origin. A major axis of variation corresponded to separation between east and west, with breeds of non-European origin concentrated in the Asian spitz and the African and Middle Eastern lineages (Figure 2C, 2D). Breeds with origins in Great Britain comprised the termini of all major western lineages, indicative of the intense selective pressure that defined Victorian-era breed development (Figure 2C) (Worboys et al., 2018). We ascertained genomic correlates of breed history potentially driving this phenomenon by estimating the correlation between maximum pseudotime value on any lineage and both homozygosity and inbreeding (F coefficient), showing that pseudotime is not significantly correlated with increased homozygosity (p=0.071l; Figure S2C, top). We did identify a significant but weak correlation with inbreeding (F coefficient; R=0.27, p=4.5E-6), likely driven by the largely purebred as opposed to mixed-breed composition of lineage termini (Figure S2C, bottom). Together, these findings suggest that identified lineages represent groups of dogs possessing similar ancestry, but which heavily diverged from wolves and ancestral canines through a combination of strong genetic drift or selection.

We then examined the distribution of modern free-breeding dogs in the PHATE embedding to determine the extent to which contemporary geographic isolation underlies trajectory structure. As expected for free-breeding dogs, geographic location was the major driving variable (Figure 2D). While contemporary purebred dogs are generally isolated, both temporally and geographically, we observed an overlap between free-breeding and purebred populations historically linked to the same geographic region (Figure 2D). This finding may reflect shared ancestry and historical admixture between these groups, likely predating formal breed formation. We also observed an accurate reflection of pseudotemporal relationships in hybridized breeds with minimal shared ancestry (Figure S2D) and in complex scenarios of mixed ancestry, such as genetic relationships between terriers, bull-type terriers, and mastiffs (Figure 1B, Figure S1D) (Fogle, 1995). Together, our findings establish a means by which global features of dog breed diversification, such as behavioral traits, can be interrogated.

### Behavioral traits are major phenotypic correlates of canine lineages

To identify global behavioral correlates of canine lineages, we generated breed-average behavioral metrics based on data from over 46,000 purebred dogs using the Canine Behavioral Assessment and Research Questionnaire (C-BARQ) (Hsu and Serpell, 2003; Serpell and Hsu, 2005). C-BARQ is a comprehensive and well-validated questionnaire for behavioral assessment comprised of owner-reported numeric scores for 100 questions related to both breed group-typical and atypical or undesirable canine behaviors (listed in Figure 3A, Methods, Table S4). We calculated 14 numeric phenotype scores per purebred C-BARQ dog, each corresponding to a single behavioral factor (Methods) (Hsu and Serpell, 2003; Serpell and Hsu, 2005).

**Figure 3.**
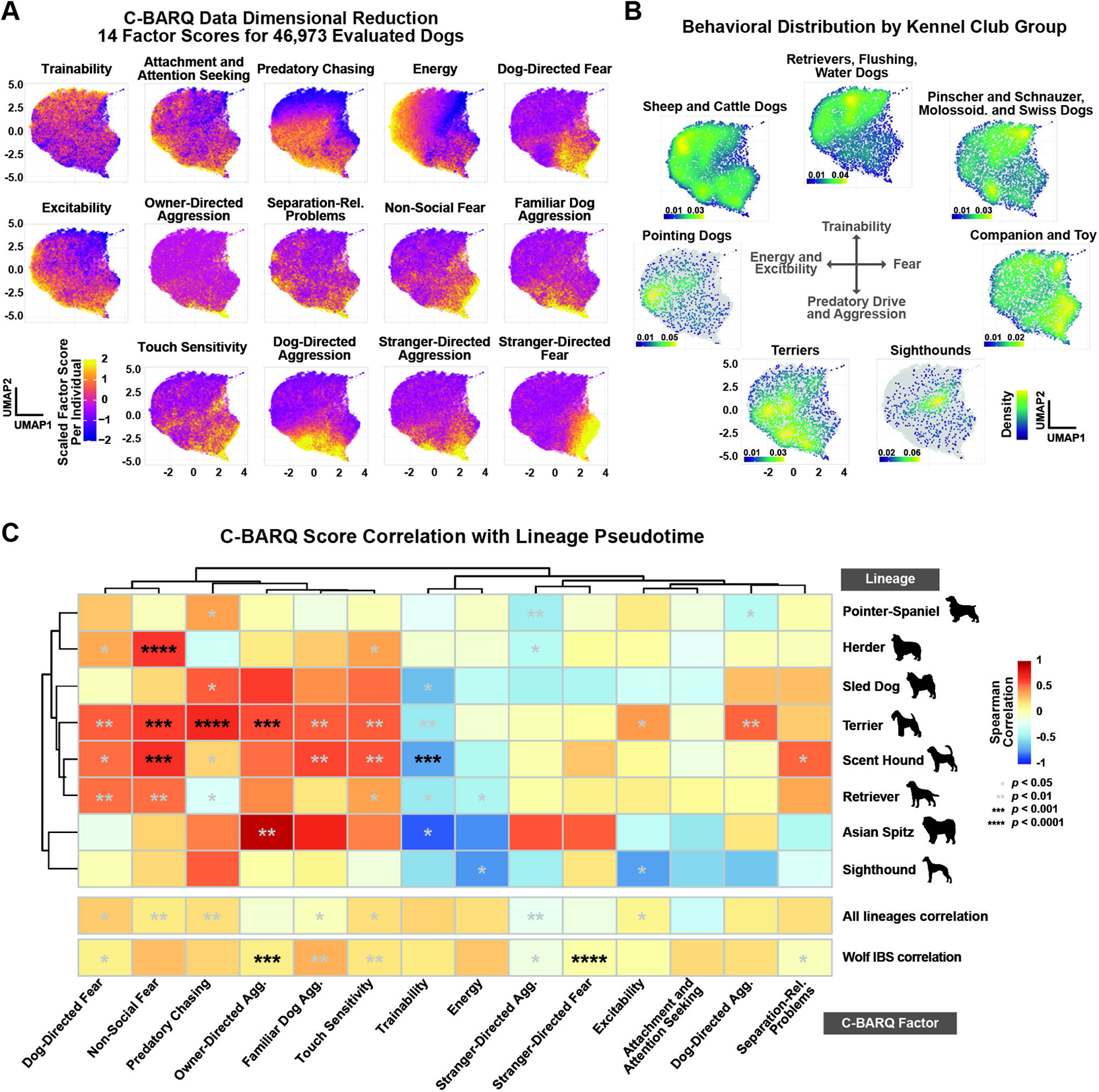
C-BARQ behavioral traits and lineage correlations. **(A)** UMAP plot generated from 14 C-BARQ factor scores for all 46,973 C-BARQ-evaluated dogs (Methods). Points indicate individual dogs and are colored by one of 14 factor scores per plot. Scores were scaled and centered (z-score) across all individuals for each factor independently. **(B)** UMAP plot (as in A) showing density of individuals within kennel club breed groups (Methods). **(C)** Heatmap showing the correlation (two-sided Spearman’s rank correlation rho) between breed-average C-BARQ factor scores and breed-average pseudotime values, maximum lineage pseudotime value (as in Figure 2A), and average gray wolf IBS (as in Figure 2B). Asterisks indicate corresponding significance values. See Figure S3C for scatterplots of selected factors and lineages.

We first verified that breeds with similar working roles broadly clustered based on calculated C-BARQ behavioral metrics. Using UMAP to dimensionally reduce the data (McInnes et al., 2018), we visualized clustering of behaviorally similar individuals and identified the major axes of behavioral variation across the dataset, which generally corresponded to fear, predatory drive and aggression, energy and excitability, and trainability (Figure 3A). We also examined the distribution of kennel club groups within the embedding, establishing that C-BARQ captures behavioral tendencies among breeds with shared historical working roles (Figure 3B, Table S4). For example, terriers largely clustered in the bottom left of the plot, consistent with predatory behavior and dog-directed aggression, while companion and toy dogs clustered along the right, consistent with increased social and non-social fear (Figure 3B). Consistent with increased trainability and decreased predatory drive and aggression, sheepdogs and retrievers clustered at the top (Figure 3B). We calculated mean C-BARQ scores for every breed in order to assign behavioral metrics to all breeds in the genetic dataset, thus reducing phenotypic noise caused by intra-breed variation and ensuring associations were not driven by a preponderance of behavioral outliers among breeds (Dodman et al., 2016; Tang et al., 2014).

To broadly characterize breed-typical behavioral phenotypes across our purebred genetic dataset, we visualized breed-average C-BARQ scores on the PHATE embedding and observed distinct suites of behavioral traits across lineages (Figure S3A). We also observed significant overlap in behavioral tendencies across multiple lineages (e.g., trainability was generally increased among herder, pointer-spaniel, and retriever trajectory breeds) (Figure S3A). Moreover, we determined that increased trainability was broadly a feature of working and sporting breeds by comparing trainability scores for working trial (e.g., border collie) versus non-working trial breeds (ANOVA p adj.=3.06E-4) and non-recognized breeds (p adj.=8.27E-3; Figure S3B, Methods).

To ascertain the degree to which genetic divergence along lineages corresponded to increasing behavioral distinctiveness, we measured the correlation between breed-average C-BARQ scores and breed-average lineage pseudotime values. We identified significant correlations for all eight lineages analyzed (dingo and African and Middle Eastern lineages excluded due to a lack of sufficient C-BARQ representation; Figure 3C, Table S5). The most significant positive correlations were between the herder lineage and non-social fear (p=1.86e-5, r=0.64), and the terrier lineage and predatory chasing (p=1.96e-5, r=0.69), the latter being consistent with working roles involving catching and killing prey (Figure 3C, Figure S3C, Table S4, Table S5).

Strikingly, the most significant negative correlation was between the scent hound lineage and trainability, consistent with selection for traits advantageous to an independently driven working style focused on following instincts rather than seeking out human cues (p=4.50e-4, r=-0.68; Figure 3C, Figure S3C, Table S4; Table S5) (Serpell and Hsu, 2005). Our results revealed a unique repertoire of behavioral correlations for each lineage consistent with working role (See Table S4); however, we also identified subsets of parallel behavior tendencies across lineages (Figure 3C, Table S4, Table S5), underscoring the utility of identifying phenotypic and genotypic correlates within lineages versus across all breeds.

### Lineage-associated variants are largely non-coding regions implicated in neurodevelopment

We next identified genetic drivers of global phenotypic shifts by performing a genome wide association study (GWAS) on the WGS subset of the described dataset (n=1,171). Using pseudotime values as quantitative phenotypes, we utilized a generalized linear model association test for the seven lineages sufficiently represented in the WGS data (See Figure 4A, top left). We identified the top 100 associated loci per lineage, ranked by index variant significance, showing that >85% were unique to a single lineage (Figure 4A and S4A). To test if the most significant associations were relevant to each lineage as a whole, rather than driven by rare variants specific to a few terminal breeds, we examined the number of breeds with non-zero lineage-associated allele frequencies, observing that such variants were generally present in >5 breeds per lineage (Figure S4B). Notably, while allele frequency distributions varied, associated variants were not generally lineage-specific but rather present at moderate allele frequencies across the entire dataset, suggesting that breed diversification was principally driven by variation that existed prior to modern breed formation (Figure S4C).

**Figure 4.**
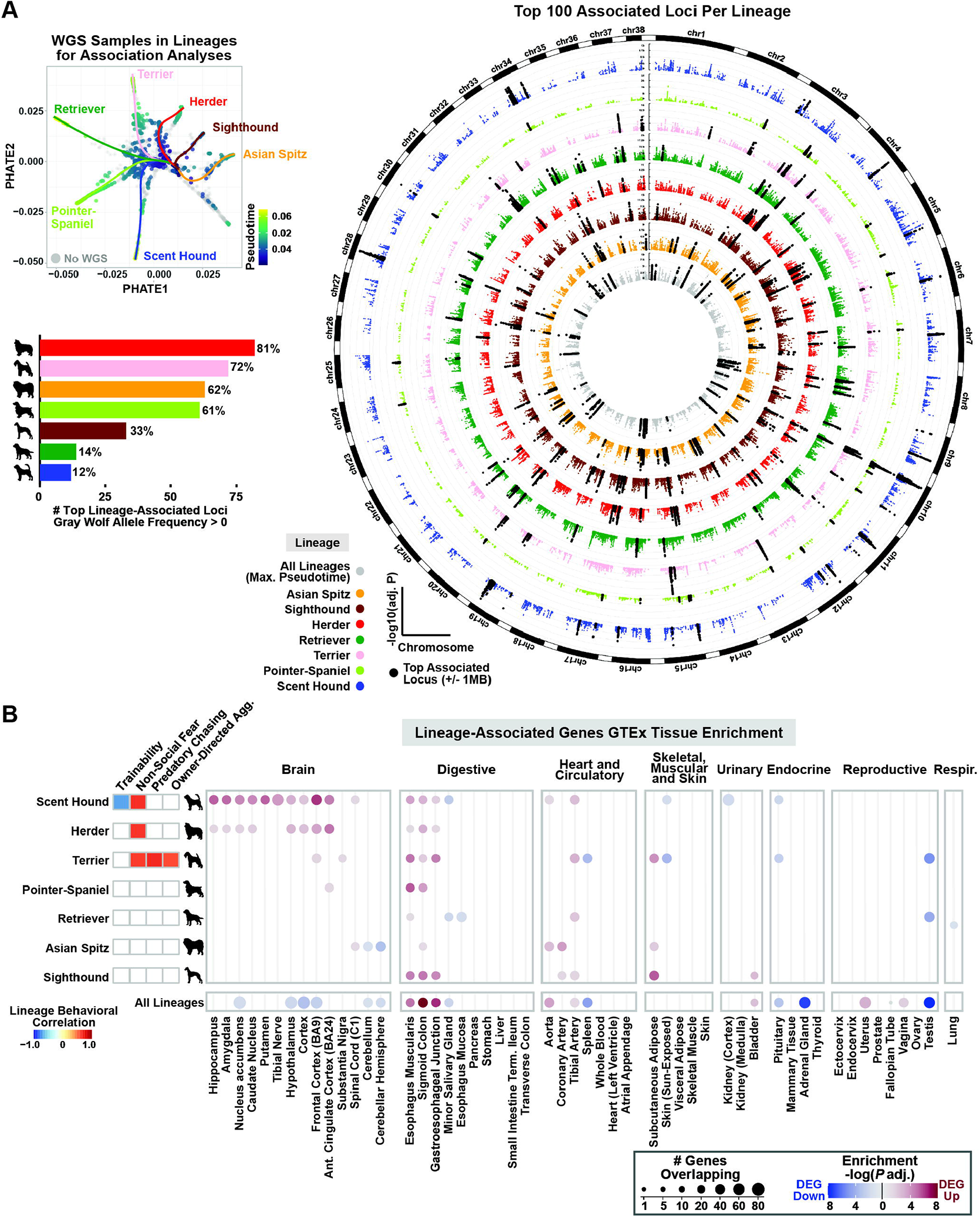
Identification and characterization of lineage-associated loci and genes. **(A)** Top Left: PHATE plot showing maximum lineage pseudotime for WGS samples only. Colored lines indicate principal curves for the seven lineages with sufficient WGS representation for association analyses (Methods). Right: Circular Manhattan plots showing log-transformed Bonferroni-adjusted GWAS p-values. Loci highlighted in black are the 100 most significantly associated per lineage. Highlighted loci were expanded to 1MB for clarity. Bottom left: Bar plot showing the number of lineage-associated alleles with gray wolf allele frequencies >0 for each of the top index variants per lineage (percentage of top loci denoted above bars). **(B)** Left: Correlation values for associations identified in the C-BARQ analysis with p<0.001 (See Figure 3C). Right: FDR-corrected significance values for tissue enrichment analysis with the GTEx v8 dataset using predicted target genes for the top 100 loci per lineage or combined lineages (Methods). Cell lines (2 of 54 GTEx samples) not shown. Circle size denotes number of differentially expressed genes (continuous scale) and circles are colored by significance (log-transformed FDR-adjusted p-values), with up and down-regulated differentially expressed genes (DEG) indicated by red and blue color scale, respectively.

To assess the degree to which this variation predates domestication, we examined allele frequencies for the top lineage-associated variants in gray wolves. For the terrier, herder, Asian spitz, and pointer-setter lineages, most associated variants were present in modern gray wolves, with 81% of the top herder lineage-associated loci present at non-zero allele frequencies, indicating that breed diversification along this lineage primarily involved selection on ancestral variation, suggestive of ancient origins for variants driving herding dog behavioral tendencies (Figure 4A, bottom left). Moreover, herder lineage-associated allele frequencies in wolves were significantly different than all but the terrier and Asian spitz lineages (p adj.=1), which had the second and third highest percentages of top loci present in gray wolves (72% and 62%, respectively; Figure 4A, bottom left). By comparison, most variants associated with the sighthound, retriever, and scent hound lineages were dog-specific, and therefore likely derived.

Characterization of lineage associations at the variant rather than locus level required assessing the degree to which variant significance predicts functionality; therefore, we applied two methods for assessing coding and non-coding functionality and determined that both base-wise conservation scores (phyloP) and predicted non-coding functional significance (DeepSEA) were skewed toward the top ranked variant at each locus (Figure S4D) (Pollard et al., 2010; Zhou and Troyanskaya, 2015). We therefore expanded our characterization of individual variants to the top five significant variants per locus and found that 74% of the top 100 loci per test contained only single nucleotide variants, and 96% contained no indels >4 bp (Figure S4E). Finally, only 1% of loci contained a non-synonymous coding variant; all missense, splice, and UTR variants were predicted to be of either low or moderate impact (Figure S4E; Methods). Strikingly, of the 16,250 significantly associated variants from all analyzed lineages, only 76 were coding variants predicted to be of moderate or high impact by either tool, and only two were predicted to be of high impact by both metrics (Table S6).

Given the apparent role of regulatory variation in driving lineage associations, we examined overlap between lineage-associated variants and loci with known enhancer and promoter functions in other species using the ENCODE database of candidate cis-regulatory elements (cCREs) (Moore et al., 2020). We found that within 95 lineage-associated loci, at least one of the top five variants overlapped a replicating human or mouse cCRE in the orthologous region (Figure S4F; Table S6). Notably, the adult and embryonic brain were the top human and mouse tissues, respectively, with the greatest percentage being human embryonic brain-specific (31%) followed by human adult brain-specific (26%) (Figure S4F; Table S6). The intersection between lineage-associated variation and regulatory elements across species indicates that a subset of lineage-associated variants may affect evolutionarily conserved regulatory elements, particularly those with tissue-specific neurodevelopmental functions.

To test if putative target genes of lineage-associated regulatory regions were enriched for certain functions, we mapped predicted gene interactions for non-coding lineage-associated loci and performed gene-based tissue expression enrichment (Methods). Using the set of genes predicted to interact with each set of lineage-associated non-coding loci (lineage-associated genes), we tested 52 human tissue types for expression enrichment (Methods) (Consortium et al., 2020). Of the 16 significantly enriched tissues for the scent hound lineage, 10 were brain tissues, as were eight of 11 from the herder lineage (Figure 4B, Table S7). Notably, the most significant scent hound and herder lineage enrichments, respectively, were the frontal cortex Brodmann area 9 and the anterior cingulate cortex Brodmann area 24, both broadly implicated in socio-cognitive functions, including learned fear response (Figure 4B, Table S7) (Apps et al., 2016; Jhang et al., 2018; Snow, 2016).

We also observed that behavioral selection was not uniform across lineages. For example, the herder, scent hound, and terrier lineages showed the most robust behavioral correlations (p<1E-3), as well as significant lineage-associated gene enrichment in one or more prefrontal areas of the cortex associated with human social cognition, decision-making, and emotional intelligence (Figure 4B) (Duncan and Owen, 2000; Fleck et al., 2006; Lane et al., 1997). However, the sighthound lineage showed no robust behavioral correlations (p>0.01), nor forebrain or midbrain enrichment for lineage-associated genes. Instead, the most significant enrichment was for subcutaneous adipose (p adj.=3.68E-6), consistent with selective breeding aimed at morphology (Figure 4B). We also verified that the predominance of brain associations among lineage-associated loci was not a general feature of canine complex traits by performing GWAS with discrete phenotypes and demographic features followed by gene-based tissue expression enrichment analyses (Figure S5, Table S8; Methods).

While identification of canine brain regions homologous to higher-order human cortical regions remains difficult, previous studies identifying significant variation in brain morphology across domestic dog breeds suggest that a subset of the lineage-associated loci identified herein may contribute to gross neuroanatomical phenotypes relevant to behavior and with neurodevelopmental origins (Hecht et al., 2019). Therefore, we also tested if expression of lineage-associated genes was enriched during specific neurodevelopmental time periods by performing gene set enrichment analysis using published expression data for 29 ages of human brain samples (Kang et al., 2011). Interestingly, the top two enrichments for any lineage (corresponding to the scent hound and pointer-setter lineages) were both at the 26 post-conception week (pcw) stage, a period during human late fetal neurodevelopment characterized by thalamocortical innervation (p adj.=6.09E-5 and p adj.=2.82E-6, respectively; See Table S7 for all lineage enrichment results) (Vasung et al., 2010). Together, these data suggest that scent hound-associated variation may principally affect innervation of the frontal cortex, evoking the hypothesis that human-directed selective pressure shaped the neurodevelopment of scent hound breeds toward a unique mode of sensory-associative learning (Alcaraz et al., 2018; Turner and Parkes, 2020).

Substantial temporal enrichment of lineage-associated genes suggested a complex mode of canine behavioral trait specification involving iterative modulation of gene co-expression networks. Therefore, we examined expression of lineage-associated genes across human neurodevelopmental stages (Kang et al., 2011). We detected co-expressed clusters of lineage-associated genes, including the scent hound lineage-associated late fetal co-expression timepoint (Figure 5). Gene ontology enrichment based on all lineage-associated genes together showed that “positive regulation of developmental process” and “positive regulation of cell differentiation” (both p adj.=7.64E-7) were the most significantly enriched processes, with 48% of genes in either implicated in “neurogenesis” (p adj.=1.16E-3), indicating a broad role for fetal neuron formation in canine behavior (Table S7).

**Figure 5.**
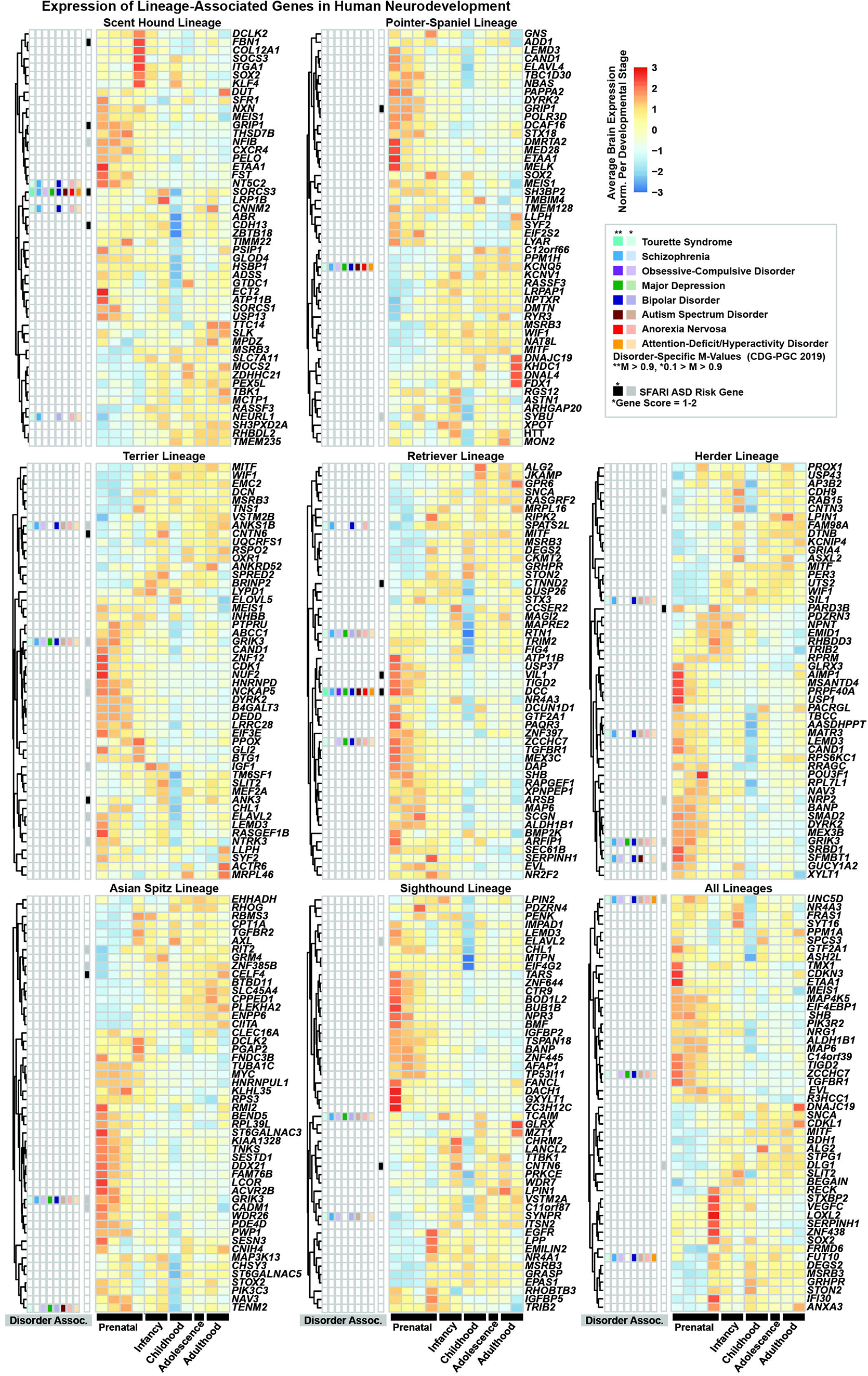
Neurodevelopmental clustering of lineage-associated genes. Heatmaps showing the average expression of the 50 most significantly lineage-associated brain-expressed genes within each of 11 human neurodevelopmental stages (Methods) (Kang et al., 2011). Heatmap values are scaled in rows. Gene order is determined by hierarchical clustering. Genes included have log2-transformed expression values >1 for at least one developmental stage (prior to row-wise scaling). Human psychiatric disorder annotations for each gene are plotted to the left. The first eight columns include annotations for eight psychiatric disorders (Methods). Colors correspond to each disorder with darker shades indicating a disorder-specific M-value (posterior probability of association) >0.9 (Consortium, 2019). Lighter shades indicate 0.1<M-value< 0.9. For SFARI ASD risk genes (black and gray squares), color coding indicates gene score according to SFARI database (Banerjee-Basu and Packer, 2010).

Given that all lineages showed associations with genes implicated in fetal brain development, and the relevance of such pathways in common human neuropsychiatric conditions, we examined the overlap between lineage-associated genes and risk genes identified in both a meta-analysis of eight DSM-defined psychiatric disorders and a database of Autism Spectrum Disorder (ASD) risk genes (Figure 5; Methods) (Banerjee-Basu and Packer, 2010; Consortium, 2019; Willsey et al., 2013). Forty-three of the top lineage-associated brain-expressed genes are identified as risk-associated in one or both datasets (Figure 5, Table S7). We also found significant enrichment of genes associated with “general risk tolerance” in humans within three lineages (pointer-setter p adj.=1.78E-4; terrier p adj.=1.33E-4; retriever p adj.=6.94E-5; Table S7) (Karlsson Linnér et al., 2019; MacArthur et al., 2017; Watanabe et al., 2017). The significant overlap between lineage-associated genes and those implicated in human behavior and psychiatric disorders suggests shared cellular mechanisms underlying neurodiversity among humans and other mammals.

### Sheepdog-associated loci are enriched near axon guidance genes

Livestock herding dogs display one of the most easily defined breed group-typical behaviors, characterized by an instinctive herding drive coupled with unique motor patterns that move herds in complex ways (Coppinger and Coppinger, 2014; Serpell, 2017; Wilcox and Walkowicz, 1989). Because the terminus of the herder lineage formed a distinct cluster of sheepdog breeds, (Figure 6A, Figure S6A), we performed a WGS-based GWAS followed by gene set enrichment analysis for this cluster, and identified a single significantly enriched KEGG pathway, “axon guidance” (hsa04360), which included 14 sheepdog cluster-associated genes (enrichment p adj.=4.25e-2; Figure 6B, Table S7, Table S9) (Liberzon et al., 2011; Watanabe et al., 2017). Eight of these genes were axon guidance receptors, four were guidance cues, and two were GTPase activating proteins (Figure 6C). We also determined that sheepdog-associated loci are significantly overrepresented near genes significantly enriched for late mid-prenatal human brain expression (p adj.=4.28E-2; Table S7). These genes converge spatiotemporally in several developmental contexts, notably in midline patterning, evoking intriguing hypotheses regarding binocular vision and motor behavior given their functional convergence in axon sorting at the optic chiasm (Figure 6C) (Ducuing et al., 2019; Larsson, 2015; Prieur and Rebsam, 2017).

**Figure 6.**
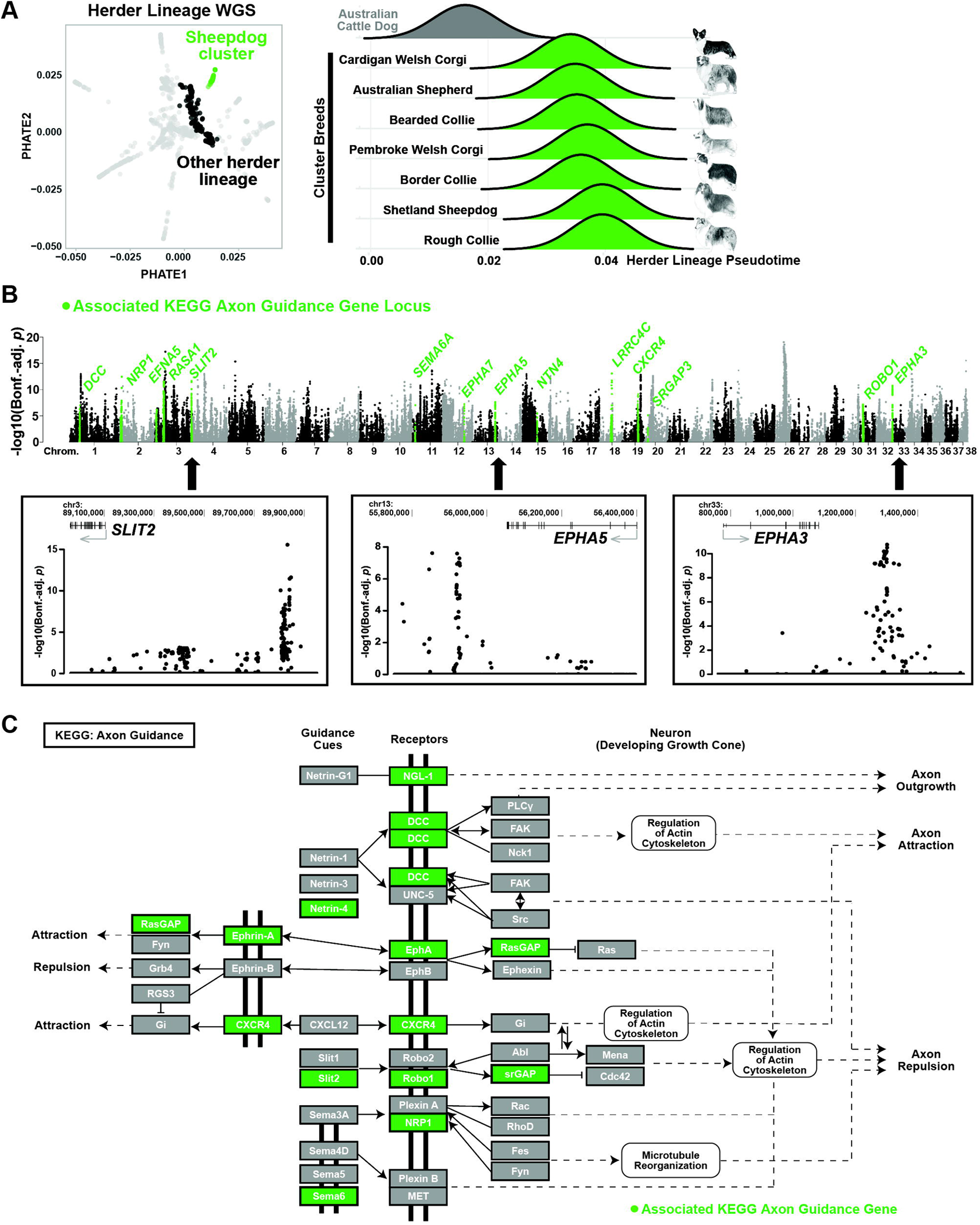
Sheepdog-associated genes are enriched for axon guidance functions. **(A)** Left: PHATE plot showing individuals with WGS data. Those with herder lineage pseudotime values >0.03 (sheepdog cluster) are green and all others with non-zero herder lineage pseudotime values (comparison group for GWAS) are black. WGS samples not part of sheepdog GWAS (non-herder lineage) are gray. Right: Ridge plot showing distribution of herder lineage pseudotime values for WGS samples within each breed in the sheepdog cluster (green) with the y-axis indicating density. The next closest breed by breed-average pseudotime values (Australian cattle dog) is also shown (gray). **(B)** Top: Manhattan plot showing sheepdog cluster GWAS results with log-transformed Bonferroni-adjusted p-values. Associated KEGG axon guidance gene loci are indicated in green. Highlighted loci were expanded to 1MB for clarity. See Methods for description of chromosome 26 peak. Bottom: genome browser images of three significantly associated loci. **(C)** Axon guidance pathway generated by KEGG (I04360) and modified for clarity with highlighted factors indicating significant gene associations.

While the associated axon guidance genes included those in netrin, slit, and semaphorin signaling pathways, the greatest number were part of the ephrin signaling pathway, including three ephrin receptor genes (Figure 6B, Figure 6C). For example, we identified an 11.3 kb sheepdog-associated locus within the same topological domain as the axon guidance gene *EPHA5*, which encodes an ephrin receptor broadly expressed in the developing brain and implicated in dopaminergic neuronal projection guidance (Figure 6B, Figure S6C) (Cooper et al., 2009; Deschamps et al., 2009). The associated haplotype of 18 variants (index SNP adj. p=2.577E-8) was present at an allele frequency of 77.5% in all border collies (83.3% in working lines; Figure S6B, Methods), 3.1% in non-sheepdog cluster herder lineage dogs, and 7.6% in all non-border collie breeds (Figure S6C). Moreover, prevalence of a core subset of variants within the border collie haplotype in wild canids indicates that human-directed pressure to produce dogs that herd sheep may have involved co-opting an ancient regulatory haplotype (Figure S6C). The striking frequency of loci associated with synaptic components may be pertinent to numerous processes relevant to behavior. For example, *EPHA5* and its ligand *EFNA5*, both sheepdog-associated genes, have been implicated in anxiety and maternal pup-gathering behavior in mouse models, suggesting that herding drive may involve augmentation of the same anxiety-associated pathways driving maternal protective behaviors in other species (Mamiya et al., 2008; Sheleg et al., 2017; Weber and Olsson, 2008). Together, our results indicate a relatively complex mode of inheritance for sheepdog-associated phenotypes involving a set of variants that iteratively modulate neuronal circuity, evoking an intriguing hypothesis regarding the ancient underpinnings of herding in mammalian behavior and the origins of genetic variants conferring such behaviors among early dogs.

## Discussion

Humans selectively bred dogs for specific occupations for thousands of years, producing a powerful system for understanding how behavior is encoded in genomes. The heritability of canine behavioral tendencies has previously been noted (MacLean et al., 2019; Svartberg, 2006); however, identification of contributing loci has historically been challenging due to the inherent complexity of canine population dynamics, featuring varying degrees of selective pressure and population bottlenecks (van Rooy et al., 2014). Embracing the intrinsically high-dimensional nature of canine genetic data while accurately distilling to its most fundamental axes of variation enabled our approach to studying canine behaviors centered on traits advantageous for distinct roles in human society. Our findings significantly advance the field of canine genetics in three major realms: we provide a substantially improved model for genetic relationships among modern dogs; reveal the biological basis of canine behavioral specification; and uncover precise molecular pathways iteratively modified to produce ideal canine workers and companions.

Our work establishes the fundamental axes by which domestic dogs have genetically deviated in the thousands of years since domestication which, contrary to recent reports, underscores the central role that behavior has played in the trajectory of canine genetic diversification (Morrill et al., 2022). One reason for the apparent discrepancy is that the lineage-based approach applied in this study enabled the detection of allele frequency shifts characteristic of deep historical relationships relevant to behavior, as opposed to between individuals or single breeds relevant to demography. Moreover, stochastic departures from expected allele frequencies are the rule rather than the exception among modern dog genomes. Therefore, we propose that future studies utilizing purebred dog genomes should be agnostic to the nature by which such mutations arose in frequency (e.g., selection versus genetic drift), as opposed to characterizing departures from expected allele frequencies among individual loci as signatures of selection accrued over short periods of time (Akey et al., 2010; Kim et al., 2018; Lindblad-Toh et al., 2005; Sabeti et al., 2006; Schlamp et al., 2016).

Ideally, future studies addressing stereotypical canine behavior will avoid phylogeny-based characterizations of modern breed relationships and, instead, account for the nature by which modern breed creation has proceeded, often through mixing phenotypes via breed hybridization as opposed to behavioral selection producing breed or lineage-specific alleles or traits. By reframing modern purebred relationships in a way that de-emphasizes variation attributable to genetic isolation of individual breeds, we circumvent difficulties faced in designing analyses with sufficient power to identify behavioral trait associations (Ilska et al., 2017; Morrill et al., 2022). Moreover, by analyzing genomes representing diverse breeds and demographic characteristics in aggregate, we identify correlations robust to the effects that environment, training, and individual breed biases have on owner-reported behavioral metrics (Dodman et al., 2018; Morrill et al., 2022; Scott and Fuller, 2012; Wells et al., 2012).

While C-BARQ is explicitly designed to assess behavioral traits associated with temperament in dogs (e.g., consistent differences in behavior such as environmental sensitivities and reactions to both social and nonsocial stimuli), these behaviors are also fundamentally linked to dogs’ historical working roles (Hsu and Serpell, 2003; MacLean et al., 2019). While some occupation-related behaviors are largely learned, others, such as a willingness to respond to human direction or an innate drive to perform certain tasks are likely to be driven by biologically-based, temperamental predispositions. This link is what makes C-BARQ particularly useful for assessing the abilities of potential working dogs (Bray et al., 2019; Duffy and Serpell, 2012), and in this study, for identifying broad lineage-associated behavioral tendencies. For example, there are many ways to describe stereotypical “herding dog” behavior. The act of herding, performing specific motor sequences, is critical, but just as important is a strong temperamental motivation to perform difficult tasks assigned and the ability to detect and interpret minute changes in human signals and livestock body language (Coppinger and Coppinger, 2014; Serpell, 2017; Wilcox and Walkowicz, 1989). One of the most significant behavioral correlations identified herein was with the herder lineage and non-social fear, anxiety-related behavior in response to phenomena that trigger fear responses, potentially related to a hyper-awareness of surroundings. This factor was also significantly correlated with the terrier and scent hound lineages, and all three lineages are those for which acute sensitivity to specific environmental stimuli is advantageous: herding dogs are keenly aware of subtle changes in the body language of livestock, terriers of stimuli signaling hidden prey, and scent hounds track the movements of game using non-visual cues (see also Table S4).

Whereas canine aesthetic traits or extreme morphological phenotypes often involve high-impact *de novo* mutations, this study supports that canine diversification along lineages is largely driven by non-coding variants, often present at moderate frequencies in non-lineage populations, suggesting a substantially more complex mode of inheritance for canine behavioral versus non-behavioral traits. Our results indicate that variants affecting brain form and function driving behavioral modulation in dogs are likely to be of relatively small-effect, demonstrating that even in scenarios of extreme phenotypic variation, such as that between dog breeds, the pleiotropic nature of neurodevelopmental genes constrains behavioral regulation to genetic mechanisms similar to those implicated in species divergence and in domestication and simulated selection scenarios alike (Alberto et al., 2018; Haygood et al., 2010; Heyne et al., 2014; Naval-Sanchez et al., 2018; Niepoth and Bendesky, 2020; Wang et al., 2018; Won et al., 2019).

While this study indicates the apparent dominance of non-coding variation in canine behavioral diversification, functional studies based on the findings herein, specifically those capable of associating regulatory variation with tissue-specific temporal shifts in developmental gene expression, will inform our understanding of the molecular underpinnings of canid behavioral modulation. Such studies will be particularly useful for ascertaining the degree to which selective pressures aimed at pleiotropic regulators of brain form and function drove the morphological and aesthetic diversification of domestic dogs and vice versa. Such findings have important implications for understanding mechanisms by which large phenotypic shifts occur within species, the potentially outsized significance of certain cell types in such processes, and the implications of artificial selection aimed at potential master-regulators of phenotype (Chong-Morrison and Sauka-Spengler, 2021; Coppinger et al., 1987; Sánchez-Villagra et al.; Wilkins et al., 2014).

Our findings are consistent with the inefficient nature of selective pressure aimed at complex behavioral traits in comparison to readily observable or even quantifiable aesthetic or morphological phenotypes, particularly those controlled by one or a few high-impact loci (Boyko et al., 2010; Hayward et al., 2016; Karlsson et al., 2007; Plassais et al., 2019). As such, we propose a model of canine behavioral diversification that largely predates modern breed formation, in which behavioral traits conducive to lineage-associated occupations (e.g., sled-pulling, herding) arose as a result of selective breeding for certain tasks over thousands of years within relatively geographically isolated populations (e.g., a need for livestock guardians in the Near East and sled dogs in the Arctic). This scenario is also consistent with origins for certain canine behaviors hypothesized to predate domestication (e.g., among wolves, herding-like interactions with ungulates when hunting), likely driven by small-effect variants at multiple loci in gray wolves but maintained in domestic dog populations in certain parts of the world (e.g., pastoral societies) (Mech et al., 2015). In future studies, ancient genomes will provide insight into temporal and geographic origins of variants identified in this study (Bergström et al., 2022; Perri et al., 2021; Schoenebeck et al., 2021; Sinding et al., 2020). Importantly, this study focuses on behavioral diversification that postdates domestication, the nature, timing, and precise geographical origins of which remain controversial (Bergström et al., 2022; Frantz et al., 2016; Larson et al., 2012; Pendleton et al., 2018; Perri et al., 2021; Savolainen et al., 2002; Thalmann et al., 2013). Nonetheless, ongoing efforts to resolve outstanding questions in dog domestication will be critical for understanding the nature of canine behavioral diversification relative to domestication-related behavioral shifts (vonHoldt et al., 2017; Wayne and vonHoldt, 2012).

### Limitations of the Study

This study encompassed an exhaustive representation of canine genetic data, including purebred dogs subjected to intense artificial selection, those developed using more permissive breed standards or mixed breeds, and semi-feral dogs that experience little or no human-imposed selective pressure. Nonetheless, expanding representation from underrepresented geographical areas would likely introduce intermediate trajectories among lineages and provide necessary context for future studies aimed at single lineages or specific populations. Importantly, the techniques used for exploratory population genetic analyses in this study are best applied to datasets of comparable or greater complexity and size as used here, such that accurate interpretation of the relevance of trajectories is possible.

While our study establishes that canine breed diversification is principally driven by non-coding variation, we did not perform structural variant analysis and therefore do not detect potentially consequential copy number variants (Serres-Armero et al., 2021), which play significant roles in mammalian brain evolution and human neurodevelopmental disorder risk (Dennis et al., 2012; Zarrei et al., 2019). Newly available long-read reference annotated genomes (Edwards et al., 2021; Field et al., 2020; Halo et al., 2021; Jagannathan et al., 2021; Wang et al., 2021) will improve accuracy in identification of CNVs and indels, as well as variant identification in GC-rich promoter regions (Kim et al., 2021; Rhie et al., 2021).

Finally, it remains difficult to ascribe finite canine behavioral traits to individual loci in much the same way that assigning loci to risk for human psychiatric disorders is complicated by diagnostic overlap between disorders, locus heterogeneity, pleiotropy, and the effects of common variation on risk (Consortium, 2019). Moreover, disentangling behavioral phenotypes from environmental factors and human bias is challenging in any mammalian system, even in laboratory environments (Schellinck et al., 2010). Nonetheless, this study provides an essential foundation for future studies examining relationships between individual variants and behavior, demonstrating that a primary outcome of behavioral diversification among domestic dog lineages is a spectrum of diverse canine genomes that principally differ in how they regulate gene expression in the brain, thus deepening our understanding of how complex mammalian behaviors are biologically encoded.

## Acknowledgements

We thank all past and present members of the Ostrander lab, as well as Ed Giniger, Laurent Frantz, Smita Krishnaswamy, James Noonan and lab, and Angeliki Louvi for insightful comments. We thank the Dog10K consortium for sequencing of NHGRI samples. We also thank Cathryn Mellersh and Brian Davis for providing samples, as well as the many dog owners who have provided DNA for these and other studies. This work was supported by funding from the Intramural Program of the National Human Genome Research Institute (E.V.D., E.A.O.).

## Author Contributions

E.V.D: Conceptualization, Data Curation, Methodology, Software, Validation, Formal Analysis, Investigation, Data Curation, Writing – Original Draft, Writing – Review & Editing, Visualization

J.A.S.: Resources, Data Curation

E.A.O.: Organization and Direction of Study, Writing – Review & Editing, Supervision, Funding Acquisition

## Inclusion and Diversity

We support inclusive, diverse, and equitable conduct of research.

## Supplemental Figure Legends

**Figure S1. Characterization of dimensionality-reduction methodologies, related to Figure 1. (A)** PCA plot (top left), UMAP plots (bottom left), and neighbor-joining tree (right) colored by kennel club group (Table S2). Each UMAP plot was generated with varying gamma, minimum distance, and spread values as indicated on each plot. **(B)** PHATE plots colored by PCA eigenvector. PC values were scaled and centered for visualization. **(C)** Three-dimensional PHATE plot (left) colored as in A with zoomed view of the central portion of the plot (right). **(D)** PHATE plot colored by clades published in Parker et al. (2017). Triangles represent members of breeds designated as “basal” in one or more of the indicated studies and italicized labels show the locations of selected “basal” breeds (Larson et al., 2012; Parker et al., 2004; vonHoldt et al., 2010).

**Figure S2. Canine lineage identification and characterization, related to Figure 2. (A)** Pseudotime values for each lineage plotted individually in PHATE plots (left) and in concentric circular heatmaps (right) which annotate tips of the neighbor-joining tree (see also annotation of tree in Figure S1A). Pseudotime values for all plots are scaled and centered (z-score) for visualization. **(B)** PHATE plots showing mean pairwise IBS values between each individual in the plot and individuals belonging to each of nine scent hound breeds. Scaling and centering (z-score) of mean pairwise IBS values was performed on the entire dataset simultaneously. All individuals belonging to the comparison breed are shown in black and their locations are highlighted with arrows corresponding to the breed centroid. Number of individuals in breed and breed-average scent hound lineage pseudotime (PT) are also shown for each breed. **(C)** Breed-average length of runs of homozygosity (kb; top) and method-of-moments F coefficient estimate (bottom) versus breed-average maximum lineage pseudotime values with associated spearman correlation rho and p-values. **(D)** PHATE plots indicating individuals belonging to selected breeds created by hybridization of existing breeds (purple), with contributing breeds (red and blue) indicated on each plot. Contributing breeds are approximated based on anecdotal breed history and/or prior genetic analyses (Parker et al., 2017; Wilcox and Walkowicz, 1989).

**Figure S3. Characterization of C-BARQ scores mapped to genetic dataset, related to Figure 3. (A)** PHATE plots showing purebred dogs colored by their respective breed-average C-BARQ score. All scores were scaled and centered for visualization. Non-purebred dogs or purebred dogs not represented in C-BARQ are colored in gray. **(B)** Left: FCI-recognized purebred dogs in PHATE plot colored by FCI working trial requirement (required in red, optional in purple, no trial in gray). Right: Boxplot showing breed-average trainability scores for non-FCI-recognized breeds, FCI-recognized breeds with no working trial requirement, an optional trial requirement, and a required trial. Significance calculations were obtained using a one-way ANOVA and Tukey test for all comparisons (Methods). **(D)** Breed-average C-BARQ score versus breed-average lineage pseudotime for the top significantly correlated factors. Significance and correlation values correspond to Spearman correlations as shown in Figure 3C. Linear regression line with 95% confidence interval is shown.

**Figure S4. Characterization of lineage-associated variants, related to Figure 4. (A)** Heatmap showing overlap between the top 100 loci for each lineage. Number of overlaps was calculated using pairwise comparisons between each lineage to identify overlap between any portion of the locus. Number of unique loci for each lineage designated in each column that overlap with lineage designated in the row name are shown. Numbers in gray squares on the diagonal show the number of loci in the corresponding lineage that have no overlap with any other lineage. **(B)** Ridge plot showing the number of breeds (within the corresponding lineage only) with non-zero minor allele frequencies for the top 100 most significantly associated index variants per lineage. **(C)** Non-lineage versus within-lineage allele frequencies for index variants for each of the top 100 loci per lineage. Color indicates locus rank (Methods). **(D)** Top: Basewise phyloP conservation scores for the top 20 variants per top 100 loci for each of seven lineages (as in panels A-C) and combined lineages by significance-based variant rank within locus. Positive scores are plotted in blue (indicating conservation). Bottom: Predicted functional significance of ranked variants. Plotted values are log-transformed predicted functional significance scores (Methods). Horizontal dashed line indicates the significance cutoff (0.05). **(E)** Left: The top 100 loci per lineage or combined lineages annotated based on the maximum insertion or deletion size within the top five most significantly associated variants per locus. Right: The number of top 100 loci per lineage or combined lineages with one of the top 5 most significantly associated variants per locus annotated as non-synonymous coding based on SnpEff (Methods). **(F)** Upset plot showing the number of lineage-associated loci (one or more of top 5 variants within the top 100 loci for any lineage) intersecting one or more human or mouse ENCODE candidate cis-regulatory elements (cCRE) by tissue after lifting the variant position to the respective genome (Methods).

**Figure S5. Comparison of lineage associations to aesthetic and morphological trait associations, related to Figure 4. (A)** Examples of Manhattan plots for lineage pseudotime associations as well as a subset of aesthetic and morphological traits and demographic features. Asterisks indicate binary phenotypes. For clarity, all log-transformed Bonferroni-adjusted GWAS p-values >0.05 are not shown. For pseudotime associations, loci highlighted in black are the 100 most significantly associated per lineage and highlighted loci were expanded to 1MB for visualization. Loci highlighted in gray boxes are lineage-associated loci overlapping selected previously identified aesthetic or morphological trait associations (Table S8) (Cadieu et al., 2009; Hayward et al., 2016; Karlsson et al., 2007; Rimbault et al., 2013; Sutter et al., 2007; Vaysse et al., 2011; Webster et al., 2015). **(B)** Bar plots showing FDR-corrected significance values for tissue enrichment analysis with the GTEx v8 dataset using predicted target genes for the top 100 associated loci for all lineages, complex traits, and demographic features. Bars are overlapping (as opposed to stacked) and colored by specific lineage or trait. Only bars representing adjusted p-values <0.05 for upregulation (indicated by horizontal dashed lines) are shown. All non-cell line GTEx v8 tissues were analyzed (as in Figure 4B), however, all tissues with no significant enrichments were omitted from plots for clarity. Asterisks indicate binary phenotypes. Longevity and standard breed height values are from Plassais et al. (2019).

**Figure S6. Characterization of the *EPHA5* sheepdog-associated locus, related to Figure 6. (A)** PHATE plot showing all purebred dogs with selected breeds within the herder lineage highlighted by anecdotal working group (Table S1). **(B)** Heatmap showing pairwise IBS values for border collies in the WGS dataset including owner report-based annotations of individuals belonging to working versus show lines. **(C)** UCSC genome browser image of a 1.4 MB locus (CanFam3 chr13:55,248,790-56,647,365) that includes sheepdog cluster-associated loci (green bars), an overlapping human neural progenitor cell (hNPC) topologically associating domain (TAD; Methods), and *EPHA5* (CanFam3 protein-coding Ensembl v104). Bonferroni-adjusted p-values for all variants are shown below bars indicating associations. Allele frequency tracks are overlayed for multiple comparisons: sheepdog cluster individuals (green; n=63) versus other non-cluster individuals in the herder lineage (black; n=321) and non-herder lineage (gray; n=787), border collies (purple; n=20) versus other individuals in the sheepdog cluster (orange; n=43), working border collies (red; n=15) versus show border collies (blue; n=5), and gray wolf (black; n=27). For the gray wolf, only allele frequencies for variants present at >75% frequency in working border collies and <25% in the entire dataset are shown for clarity. Frequency tracks for the red wolf, coyote, golden jackal, dhole, and Andean fox (black) indicate positions at which the individual is homozygous for the aforementioned subset of variants.

## RESOURCE AVAILABILITY

### Lead contact

Further information and requests for resources and reagents should be directed to and will be fulfilled by the lead contact, Elaine A. Ostrander. For specific information regarding C-BARQ data, please contact James Serpell.

### Materials availability statement

This study did not generate new unique reagents.

### Data and code availability

- Whole genome sequencing (WGS) data have been deposited at NCBI Sequence Read Archive (SRA) and are publicly available as of the date of publication. Accession numbers are listed in the Key Resources Table and Table S1. SNP array data have been deposited at Gene Expression Omnibus (GEO) and are publicly available as of the date of publication. Accession numbers are listed in the Key Resources Table and Table S1. This paper also analyzes existing, publicly available WGS and SNP array data. These accession numbers are listed in the Key Resources Table and Table S1.
- This paper does not report original code.
- Any additional information required to reanalyze the data reported in this paper is available from the lead contact(s), with access to C-BARQ data available from James Serpell upon request.

## METHOD DETAILS

### Data Ascertainment

We compiled a genomic dataset of 1,171 canid whole genome sequences obtained from the Sequence Read Archive (SRA; http://www.ncbi.nlm.nih.gov/sra) or generated by the NIH Intramural Sequencing Center. Accession and biosample numbers for previously published and unpublished WGS samples are listed in Table S1. Of the publicly available sequences used herein, 364 were provided by NHGRI and sequenced by the Dog 10K Consortium (PRJNA648123) (Ostrander et al., 2019), 166 were produced by the University of Missouri, 152 were generated by the Institute of Genetics, University of Bern, Switzerland, and 260 sequences were previously produced by the National Institutes of Health. For new WGS data, read alignment to the CanFam 3.1 reference genome (Hoeppner et al., 2014; Lindblad-Toh et al., 2005) and variant calling was performed based on GATK guidelines and as previously described (Plassais et al., 2019; Van der Auwera et al., 2013) using BWA (0.7.17), Samtools (1.6), PicardTools (2.9.2), and GATK (4.0) for BQSR. SNVs and small indels were called using GATK HaplotypeCaller (4.0.12.0) (Li and Durbin, 2009; Li et al., 2009; Van der Auwera et al., 2013).

We compiled a dataset of 3,090 Illumina Canine HD SNP chip (Illumina, San Diego, CA) arrays, 2,135 of which were previously published (Table S1). SNP array accession information for previously published and unpublished data is available in Table S1. New SNP array data was generated according to manufacturer protocols and genotype calls were made with Genome Studio V2011.1 with genotyping module v1.9.4 (Illumina).

Previously unpublished WGS and SNP array data were generated from domestic dog samples collected with owners signed consent in accordance with standard protocols approved by the NHGRI IACUC committee, protocol #GFS-05-1. Blood draws for sample ascertainment were performed by licensed veterinarians or veterinary technicians and saliva samples were collected by the owner or a member of the Ostrander lab as previously described (Plassais et al., 2019).

The combined WGS and SNP dataset (n=4,261) is comprised of 2,823 purebred dogs (FCI-recognized; owner-reported or registered), 687 non-recognized or mixed breed dogs (including three wolf-dog hybrid breeds, one of which is FCI-recognized), 658 village dogs, and 93 wild canids. Table S1 contains sample metadata including FCI breed standard-based information, and collection location for village dogs and wild canids.

### WGS and SNP Data Quality Control

The 3,090 SNP array datasets used in this study were filtered based on sample and variant missingness and include only biallelic autosomal SNVs. We extracted SNP array variants from the filtered WGS dataset (WGS filtering described below) to generate a combined SNP dataset for 4,261 individuals. The maximum sample missingness for the final dataset of 102,764 SNPs was 0.20%. Duplicate samples from both datasets combined were removed using kinship analysis with KING by calculating a cutoff value based on kinship values between known duplicates in the dataset (Manichaikul et al., 2010). Duplicate samples were pruned by keeping the sample with the greater sequencing depth for WGS data or lower missingness for SNP array data. After filtering, the maximum kinship value for the combined dataset of WGS and SNP array samples was 0.35.

WGS data was filtered to remove low quality samples and genotypes; hard filtering was performed based on GATK guidelines using bcftools: bcftools filter -e ‘FS>60.0 || SOR>3 || MQ<40 || MQRankSum<-12.5 || MQRankSum>12.5 || QD<2.0 || ReadPosRankSum<-8.0’ -Ob | bcftools filter -S. -e ‘FMT/DP<3 | FMT/GQ<10’ | bcftools filter -e ‘F_MISSING > 0.05 || MAF <= 0.0017’ -Ob | bcftools view -e ‘AC==0 || AC==AN’ -m2 -M2 (Danecek et al., 2021; Li, 2011; Van der Auwera et al., 2013). The allele frequency threshold was determined based on a requirement for at least four minor alleles in the final dataset. The filtered WGS dataset of 1,171 samples contained 20,667,447 variants with a maximum sample missingness of 6.95% and minimum sequencing depth of 10x. The average WGS sample sequencing depth and missingness after filtering were 23x and 1.48%, respectively. Inbreeding (F coefficient) and homozygosity were calculated using PLINK v1.9 (Chang et al., 2015; Purcell et al., 2007).

### Dimensionality reduction

Principal component analysis (PCA) was performed using a variance-standardized relationship matrix based on the combined dataset of 102,764 SNPs (SNP array positions) for 4,261 samples generated with PLINK v1.9 (Chang et al., 2015; Purcell et al., 2007). Percent variance explained was calculated using standard deviations of the principal components derived from prcomp (R 4.0.0 Stats package). For subsequent analyses, 20 PCs (accounting for 74.9% of the cumulative variance) were used, with subsequent PCs each accounting for less than 0.53% of variance. UMAP was run on a matrix containing the first 20 PCs using the umap package in R (R v4.1.0, umap 0.2.7.0) (McInnes et al., 2018). UMAP plots were generated with multiple gamma, minimum distance, and spread values as indicated on plots in Figure S1A. The two-dimensional PHATE embedding was generated using the first 20 PCs as input with phate(pca_matrix, ndim=2, knn=80, gamma=0, decay=100) (R v4.1.0, phateR v1.0.7, python 3.7) (Moon et al., 2019). Three-dimensional PHATE was run with the same settings but with ndim=3 (Moon et al., 2019).

### Pseudotemporal Reconstruction

Slingshot (R 3.6.2, Slingshot v1.4.0) was used for identifying lineages using PHATE coordinates as input and clusters defined using kmeans(centers=40) (R 3.6.2 Stats package) with the gray wolf, red wolf, and coyote-containing cluster set as the starting cluster (Street et al., 2018). Pseudotime values were obtained for each of the 10 Slingshot-identified lineages using slingPseudotime (Street et al., 2018) (Table S3). Pseudotime values corresponding to at least one lineage were assigned to every individual in the dataset, with 74% of purebred and mixed breed dogs belonging to a single lineage. On average, each lineage included 663 individuals from 67 breeds (range=17-125; Table S3).

### Phylogenetic Analysis

For phylogenetic analysis, linkage pruning was performed on the merged SNP dataset with PLINK v1.9 using --indep 50 5 2 (Chang et al., 2015; Purcell et al., 2007). Pruned SNPs were used for distance matrix generation with PLINK v1.9 using –distance 1-ibs square for neighbor-joining tree generation with Phylip v3.697 (100 matrices generated, neighbor and consense functions) (Chang et al., 2015; Felsenstein, 2005; Purcell et al., 2007).

### Analysis of C-BARQ behavioral data

C-BARQ survey data was compiled as of July 2021 and contained 67,970 responses. Scores for the 14 previously defined behavioral factors assessed in C-BARQ (trainability, stranger-directed aggression, owner-directed aggression, dog-directed aggression, familiar dog aggression, dog-directed fear, stranger-directed fear, nonsocial fear, touch sensitivity, separation-related problems, excitability, attachment/attention-seeking, predatory chasing, energy level) were calculated as averages of the raw response values for the corresponding questions (Duffy and Serpell, 2012; Hsu and Serpell, 2003; Serpell and Hsu, 2005). In Table S4, we enumerate hypotheses related to historical canine working roles that were generated based on results in this study, each pertaining to one of 14 C-BARQ factors. Descriptions of C-BARQ factors in Table S4 are from vetapps.vet.upenn.edu/cbarq/about.cfm. Importantly, these 14 descriptions of C-BARQ factors characterize the types of questions that were asked in order to assess each of 14 aspects of temperament; however, the associations identified herein are not with one queried metric individually (e.g., fearful or wary responses to sudden or loud noises) but rather with the constellation of behaviors associated with an aspect of temperament based on multiple questions (e.g., propensity to display non-social fear responses). Several previous studies demonstrate the reliability and validity of the C-BARQ data survey, including concordance between responses from different members of the same household and factor structure consistency across cultures (Canejo-Teixeira et al., 2018; Duffy and Serpell, 2012; Hsu and Serpell, 2003; Jakuba et al., 2013; Nagasawa et al., 2011; Serpell and Hsu, 2005).

We filtered the C-BARQ dataset to eliminate multiple entries per individual as well as individuals missing greater than 20% of responses for any one factor, resulting in a dataset of factor scores for 46,973 unique individuals. The filtered C-BARQ dataset contained 237 domestic dog breeds (as reported by owner) with an average of 136 individuals per breed. We generated a UMAP plot for all individuals based on a matrix of C-BARQ factor scores for each individual using uwot (v0.1.11) in R (v3.6.2) (McInnes et al., 2018). Density of points within UMAP plot by group was calculated as two-dimensional kernel estimations evaluated using 100 grid points in each direction (MASS v7.3-51.5).

For correlation analyses, we generated breed-average C-BARQ factor scores for the 132 breeds having at least 30 individuals in the C-BARQ dataset as well as representation in the genetic dataset. The dingo and African and Middle Eastern lineages were eliminated from correlation analyses due to lack of sufficient representation in the C-BARQ dataset. For each lineage, we calculated breed-average pseudotime values, requiring that a majority of individuals in the breed belonged to the lineage in order to exclude breed averages based on single outlier individuals. We calculated two-sided Spearman’s rank correlation rho and p-values between breed-average pseudotime value and breed-average C-BARQ factor score for each lineage independently for each of 14 behavioral factors using cor(method=”spearman”) (R 3.6.2 Stats package). We used the same method to calculate correlations with breed-average gray wolf mean IBS (mean IBS between each individual and each gray wolf calculated with PLINK v1.9) and breed-average maximum lineage pseudotime (Chang et al., 2015; Purcell et al., 2007). Working trial requirements for FCI-recognized breeds were obtained from FCI breed standards (fci.be/nomenclature/). ANOVA and Tukey tests for behavioral comparisons between working trial groups were performed with all breed averages using aov and TukeyHSD (R 3.6.2 Stats package).

### Lineage Association Analyses

Lineage pseudotime GWAS were performed using WGS data from 1,167 individuals. GWAS were performed separately for 7 lineages (Sled dog, African and Middle Eastern, and dingo lineages were eliminated due to lack of sufficient representation in the WGS dataset) and also with maximum lineage pseudotime per individual, regardless of lineage. GWAS were performed using the PLINK v2.3 association analysis command with –mac 10 –glm –adjust gc and –lamba where the lambda value was set as the LD score regression intercept calculated using LDSC (Bulik-Sullivan et al., 2015; Chang et al., 2015; Purcell et al., 2007). The LD score regression intercepts for each lineage association were calculated with ldsc.py –l2 –ld-wind-kb 1000 –maf 0.004 followed by munge_sumstats.py and ldsc.py –h2 (Bulik-Sullivan et al., 2015). Bonferroni-adjusted p-values from each GWAS were used for the variant clumping procedure implemented with PLINK (v1.9) using –maf 0.004 –clump-p2 0.01 –clump-p1 0.001 –clump-r2 0.50 –clump-kb 1000 –clump-field BONF in order to define start and end points of associated loci (Chang et al., 2015; Purcell et al., 2007). The minor allele frequency (MAF) cutoff of 0.004 corresponds with restriction to variants with at least 10 minor alleles in the WGS dataset to match the cutoff (--mac 10) implemented in the association analysis command. Coordinates of loci were calculated based on the first and last positions of clumped variants.

For analyses based on the top 100 loci per lineage, locus ranks were calculated based on the index variant Bonferroni-adjusted p-value. Of the top 100 loci for each of the seven individual lineages and combined lineage GWAS, 42% of loci were less than 10 kbp in length and the minimum and maximum Bonferroni-adjusted index SNP p-values for the top 100 loci for any lineage were 2.54E-32 and 1.22E-4, respectively (Table S6). Circular Manhattan plot was generated using CMplot 3.7.0.

Significance tests to compare distributions of wolf allele frequencies for lineage-associated variants were performed with Kolmogorov-Smirnov tests between all lineages using ks.test (R 3.6.2 Stats package) followed by Bonferroni correction. Prior to wolf allele frequency calculations and Kolmogorov-Smirnov tests, the lineage-associated allele was set as the alternate allele if present at a higher allele frequency in samples with lineage pseudotime values greater than the median versus less than the median (per lineage).

Association analyses based on morphological and aesthetic traits and demographic features were performed using PLINK v2.3 and LDSC as described above for quantitative phenotypes (height, weight, longevity) as well as binary phenotypes (white chest, drop ears, furnishing, bulky, muscled, long legs, working trial requirement, Great Britain breed origin, Italy breed origin, France breed origin) (Bulik-Sullivan et al., 2015; Chang et al., 2015; Purcell et al., 2007). Breed origin countries are based on FCI breed standards (fci.be/nomenclature/). Great Britain, Italy, or France is the FCI-based breed country of origin for 44% of the purebred individuals in the dataset and 38% of the FCI-recognized breeds in the dataset (see Figure 2C). All phenotypes were applied to purebred individuals based on breed (Table S1). All phenotypes other than standard breed weight were obtained from previously published metadata as described in Table S8 and original publications for loci highlighted in Figure S5 are listed in Table S8 (Cadieu et al., 2009; Hayward et al., 2016; Karlsson et al., 2007; Plassais et al., 2019; Plassais et al., 2017; Rimbault et al., 2013; Sutter et al., 2007; Vaysse et al., 2011; Webster et al., 2015).

### Variant annotation and enrichment tests

Variant coordinates were lifted from CanFam3.l to hg19 (GRCh37), mm10 (GRCm38), and hg38 (GRCh38) genome builds with liftOver and default settings (Rhead et al., 2010). We required that lifted variants reciprocally mapped to the correct CanFam3.1 position. DeepSEA was performed on reciprocally-lifting hg19 coordinates using rundeepsea.py (Zhou and Troyanskaya, 2015). DeepSEA functional significance scores were calculated for each canine variant that had at least one allele matching the reference human allele by running DeepSEA with both canine alleles set as the human reference allele to account for either allele matching the human reference allele (approximately 57% of reciprocally lifting variants) (Zhou and Troyanskaya, 2015). To calculate phyloP scores for each variant mapped to hg38, we used bigWigAverageOverBed and hg38.phyloP100way.bw obtained from the UCSC Genome Browser at http://hgdownload.cse.ucsc.edu (Pollard et al., 2010; Rhead et al., 2010). We used both SnpEff (v5.0 with CanFam3.1.99) and VEP (v104) to identify lineage-associated coding variants and predicted coding effects (Cingolani et al., 2012; McLaren et al., 2016).

Lifted coordinates were intersected with all cCRE regions for human (n=926,535) and mouse (n=339,815) obtained from ENCODE (Moore et al., 2020). Regulatory element annotations for tissues were obtained using these ENCODE cCRE IDs and merged for all human and mouse tissues that had replicate samples available in the SCREEN database (data for individual tissues listed in Table S6). All cCREs were treated as active if annotated as anything other than unclassified or low-DNase (any combination of H3K27ac, H3K4me3, CTCF, DNase). cCREs were filtered for those that replicate at least once within the replication groups as described in Table S6, resulting in 261,643 human cCREs and 169,594 mouse cCREs. Mouse adult brain, mouse adult liver, and mouse adult spleen were removed due to lack of available replicates. All cCREs with intersecting lineage-associated variants were also lifted to CanFam3.1 with liftOver using -minMatch=0.5 to require reciprocal mapping back to the correct CanFam3.1 position (Rhead et al., 2010).

Predicted target genes for all loci were identified using GREAT (McLean et al., 2010). First, associated locus coordinates were defined as described above. The median locus size for input to GREAT was 42.7 kb. All clumps were lifted to hg38 using liftOver with minMatch=0.5 (Rhead et al., 2010). Next, the rGREAT client for GREAT analysis R package (v1.20.0) was used to predict target genes using submitGreatJob (includeCuratedRegDoms=FALSE) and getEnrichmentTables with the default GREAT gene regulatory domain definition (Basal plus extension) (McLean et al., 2010). Genes predicted for the top 100 loci per lineage or combined lineages (172-188 genes per lineage) were used as input to FUMA (v1.3.7) for gene set enrichment, with the background gene set specified as protein coding genes (ensemble v92). Human gene expression values and enrichment significance values were obtained with FUMA based on the GTEx v8 (54 sample types, of which 52 are tissues versus cell lines; 30 general tissue types) and BrainSpan (29 different ages of brain samples and 11 general developmental stages of brain samples) datasets (Consortium et al., 2020; Kang et al., 2011; Watanabe et al., 2017). The 11 developmental stages of BrainSpan human brain tissue are as follows: early prenatal (8-12 pcw), early mid-prenatal (13-18 pcw), late mid-prenatal (19-24 pcw), late prenatal (25-38 pcw), early infancy (0-5m), late infancy (1-18m), early childhood (19m-5y), late childhood (6-11y), adolescence (12-19y), young adulthood and middle adulthood (Kang et al., 2011). All optional parameters were set to defaults (Benjamini-Hochberg multiple testing correction for gene-set enrichment testing, maximum adjusted p-value for gene set association <0.05, and minimum overlapping genes ≥ 2) (Benjamini and Hochberg, 1995; Watanabe et al., 2017). Human behavioral trait GWAS gene set enrichment was performed using FUMA and was based on the NHGRI GWAS catalog version e104 (2021-09-15) (MacArthur et al., 2017; Watanabe et al., 2017). Hierarchical clustering in all heatmaps was performed using pheatmap (Kolde, 2015)

### Sheepdog cluster GWAS and locus annotation

Sheepdog cluster GWAS, variant clumping, and locus annotation were performed as described above except with case/control (case n=63 sheepdog cluster individuals; control n=321 non-sheepdog cluster herder lineage individuals; Table S1) rather than a quantitative phenotype with PLINK v1.9 (Chang et al., 2015; Purcell et al., 2007). KEGG pathway gene set enrichment analysis was performed using FUMA v1.3.7 as described above. Owner-reported data was used to identify working border collies versus those bred for aesthetic competitions (show border collies) and designations were verified based on pairwise IBS data as shown in Figure S6B.

The broad sheepdog cluster-associated region evident on chromosome 26 (Figure 6B) is comprised of two adjacent clumps each spanning approximately 1 MB (the maximum allowed size for variant clumping, Methods). The first region is chr26:21141724-22111646, index variant p adj.=9.09E-20 (chr26:21264487), index variant-assoc. genes*: PITPNB, TTC28*, other associated genes: *MN1*. The adjacent region is chr26:22411585-23367720, index variant p adj.=2.43E-19 (chr26:22645274), index variant assoc. genes: *CABP7, NF2,* other associated genes: *KREMEN1, EMID1, RHBDD3, GAS2L1, EWSR1, RASL10A, RFPL1, NEFH, THOC5, NIPSNAP1, ZMAT5, UQCR10, ASCC2, HORMAD2, MTMR3.* Both index variants are present at 100% allele frequency in show border collies versus 26.7% and 30%, respectively, in working border collies. In addition, both index variants are present at 100% allele frequency in rough collies and at ≥70% in two other cluster breeds: Pembroke Welsh Corgi and Shetland Sheepdog. Among all non-sheepdog cluster individuals in the entire dataset, both index variants are present at <2% allele frequency.

Previously published WGS data were obtained for the golden jackal (PRJNA274504, SAMN03366713), dhole (PRJNA266585, SAMN03168405), and Andean fox (PRJNA232497,

SAMN02487034) for the purposes of identifying allele conservation in Figure S6C, but were not used in any other analyses. UCSC genome browser images show CanFam3.1 protein-coding genes (Ensembl v104) and human neural progenitor cell (hNPC) topologically associating domain (TAD) regions (Aken et al., 2016; Dixon et al., 2015; Rhead et al., 2010) and were lifted to CanFam3.1 as described above.

## QUANTIFICATION AND STATISTICAL ANALYSIS

Two-sided Spearman’s rank correlation rho and p-values between breed-average pseudotime value and breed-average C-BARQ factor score (Figure 3C, Figure S3C) were calculated using cor(method=”spearman”) (R 3.6.2 Stats package). Spearman correlation rho and p-values calculated for runs of homozygosity (kb) and inbreeding (F) versus breed-average maximum lineage pseudotime (Figure S2C) were calculated using stat_cor(method=“spearman”) in ggpubr (0.4.0). ANOVA and Tukey tests for behavioral comparisons between working trial groups (Figure S3B) were performed using aov and TukeyHSD (R 3.6.2 Stats package). Boxplots in Figure S3B were generated with ggplot2 3.3.5 with default settings, and show the median and first and third quartiles (Wickham, 2016). Bonferroni adjustment of p-values for GWAS was performed using PLINK (v1.9) (Chang et al., 2015; Purcell et al., 2007). Significance tests to compare distributions of wolf allele frequencies for lineage-associated variants were performed with Kolmogorov-Smirnov tests using ks.test followed by Bonferroni correction using p.adjust (R 3.6.2 Stats package). Benjamini-Hochberg multiple testing correction was performed for gene-set enrichment testing as implemented in FUMA v1.3.7 (Benjamini and Hochberg, 1995; Watanabe et al., 2017).

## Supplemental Tables

**Table S1. Genetic dataset metadata and accession information**, related to Figure 1 and Methods

**Table S2. Descriptions of FCI breed groups**, related to Figure 1 and Methods

**Table S3. PHATE embedding coordinates and lineage pseudotime values**, related to Figure 1 and Figure 2

**Table S4. Descriptions of C-BARQ factors and associated findings**, related to Figure 3 and Methods

**Table S5. C-BARQ correlation data**, related to Figure 3

**Table S6. Lineage association GWAS and locus annotation**, related to Figures 4 and 5

**Table S7. Gene set enrichment results**, related to Figures 5 and 6

**Table S8. Phenotype data for morphological, aesthetic, and demographic GWAS**, related to Figure S5

**Table S9. Sheepdog cluster GWAS and locus annotation**, related to Figure 6

## References

1. Aken, B.L., Ayling, S., Barrell, D., Clarke, L., Curwen, V., Fairley, S., Fernandez Banet, J., Billis, K., García Girón, C., Hourlier, T., et al. (2016). The Ensembl gene annotation system. Database 2016, baw093. 10.1093/database/baw093

2. Akey, J.M., Ruhe, A.L., Akey, D.T., Wong, A.K., Connelly, C.F., Madeoy, J., Nicholas, T.J., and Neff, M.W. (2010). Tracking footprints of artificial selection in the dog genome. Proc. Natl. Acad. Sci. USA 107, 1160–1165. 10.1073/pnas.0909918107

3. Alberto, F.J., Boyer, F., Orozco-terWengel, P., Streeter, I., Servin, B., de Villemereuil, P., Benjelloun, B., Librado, P., Biscarini, F., Colli, L., et al. (2018). Convergent genomic signatures of domestication in sheep and goats. Nat. Commun. 9, 813. 10.1038/s41467-018-03206-y

4. Alcaraz, F., Fresno, V., Marchand, A.R., Kremer, E.J., Coutureau, E., and Wolff, M. (2018). Thalamocortical and corticothalamic pathways differentially contribute to goal-directed behaviors in the rat. Elife 7, e32517. 10.7554/eLife.32517

5. Amir, E.-a.D., Davis, K.L., Tadmor, M.D., Simonds, E.F., Levine, J.H., Bendall, S.C., Shenfeld, D.K., Krishnaswamy, S., Nolan, G.P., and Pe’er, D. (2013). viSNE enables visualization of high dimensional single-cell data and reveals phenotypic heterogeneity of leukemia. Nat. Biotechnol. 31, 545–552. 10.1038/nbt.2594

6. Apps, M.A.J., Rushworth, M.F.S., and Chang, S.W.C. (2016). The anterior cingulate gyrus and cocial cognition: Tracking the motivation of others. Neuron 90, 692–707. 10.1016/j.neuron.2016.04.018

7. Banerjee-Basu, S., and Packer, A. (2010). SFARI Gene: an evolving database for the autism research community. Dis. Model Mech. 3, 133–135. 10.1242/dmm.005439

8. Bannasch, D.L., Baes, C.F., and Leeb, T. (2020). Genetic variants affecting skeletal morphology in domestic dogs. Trends Genet. 36, 598–609. 10.1016/j.tig.2020.05.005

9. Benjamini, Y., and Hochberg, Y. (1995). Controlling the false discovery rate: a practical and powerful approach to multiple testing. J. R. Stat. Soc. B 57, 289–300.

10. Bergström, A., Stanton, D.W.G., Taron, U.H., Frantz, L., Sinding, M.-H.S., Ersmark, E., Pfrengle, S., Cassatt-Johnstone, M., Lebrasseur, O., Girdland-Flink, L., et al. (2022). Grey wolf genomic history reveals a dual ancestry of dogs. Nature 607, 313–320. 10.1038/s41586-022-04824-9

11. Boyko, A.R., Quignon, P., Li, L., Schoenebeck, J.J., Degenhardt, J.D., Lohmueller, K.E., Zhao, K., Brisbin, A., Parker, H.G., vonHoldt, B.M., et al. (2010). A simple genetic architecture underlies morphological variation in dogs. PLoS Biol. 8, e1000451–e1000451. 10.1371/journal.pbio.1000451

12. Bray, E.E., Levy, K.M., Kennedy, B.S., Duffy, D.L., Serpell, J.A., and MacLean, E.L. (2019). Predictive models of assistance dog training outcomes using the Canine Behavioral Assessment and Research Questionnaire and a standardized temperament evaluation. Front. Vet. Sci. 6, 49. 10.3389/fvets.2019.00049

13. Bulik-Sullivan, B.K., Loh, P.-R., Finucane, H.K., Ripke, S., Yang, J., Patterson, N., Daly, M.J., Price, A.L., Neale, B.M., and Schizophrenia Working Group of the Psychiatric Genomics, C. (2015). LD Score regression distinguishes confounding from polygenicity in genome-wide association studies. Nat. Genet. 47, 291–295. 10.1038/ng.3211

14. Cadieu, E., Neff, M.W., Quignon, P., Walsh, K., Chase, K., Parker, H.G., Vonholdt, B.M., Rhue, A., Boyko, A., Byers, A., et al. (2009). Coat variation in the domestic dog is governed by variants in three genes. Science 326, 150–153. 10.1126/science.1177808

15. Canejo-Teixeira, R., Almiro, P.A., Serpell, J.A., Baptista, L.V., and Niza, M. (2018). Evaluation of the factor structure of the Canine Behavioural Assessment and Research Questionnaire (C-BARQ) in European Portuguese. PLoS One 13, e0209852. 10.1371/journal.pone.0209852

16. Chang, C.C., Chow, C.C., Tellier, L.C., Vattikuti, S., Purcell, S.M., and Lee, J.J. (2015). Second-generation PLINK: rising to the challenge of larger and richer datasets. GigaScience 4, 7. 10.1186/s13742-015-0047-8

17. Chong-Morrison, V., and Sauka-Spengler, T. (2021). The cranial neural crest in a multiomics era. Front. Physiol. 12, 634440. 10.3389/fphys.2021.634440

18. Cingolani, P., Platts, A., Wang, L.L., Coon, M., Nguyen, T., Wang, L., Land, S.J., Lu, X., and Ruden, D.M. (2012). A program for annotating and predicting the effects of single nucleotide polymorphisms, SnpEff: SNPs in the genome of Drosophila melanogaster strain w1118; iso-2; iso-3. Fly 6, 80–92. 10.4161/fly.19695

19. Consortium, C.-D.G.o.t.P.G. (2019). Genomic relationships, novel loci, and pleiotropic mechanisms across eight psychiatric disorders. Cell 179, 1469–1482.e1411. 10.1016/j.cell.2019.11.020

20. Consortium, G., Aguet, F., Anand, S., Ardlie Kristin, G., Gabriel, S., Getz Gad, A., Graubert, A., Hadley, K., Handsaker Robert, E., Huang Katherine, H., et al. (2020). The GTEx Consortium atlas of genetic regulatory effects across human tissues. Science 369, 1318–1330. 10.1126/science.aaz1776

21. Cooper, M.A., Kobayashi, K., and Zhou, R. (2009). Ephrin-A5 regulates the formation of the ascending midbrain dopaminergic pathways. Dev. Neurobiol. 69, 36–46. 10.1002/dneu.20685

22. Coppinger, L., and Coppinger, R. (2014). Dogs for herding and guarding livestock. Livestock Handling and Transport: Fourth Edition, 245–260.

23. Coppinger, R., Glendinning, J., Torop, E., Matthay, C., Sutherland, M., and Smith, C. (1987). Degree of behavioral neoteny differentiates canid polymorphs. Ethology 75, 89–108.

24. Danecek, P., Bonfield, J.K., Liddle, J., Marshall, J., Ohan, V., Pollard, M.O., Whitwham, A., Keane, T., McCarthy, S.A., Davies, R.M., et al. (2021). Twelve years of SAMtools and BCFtools. GigaScience 10, giab008. 10.1093/gigascience/giab008

25. Dennis, M.Y., Nuttle, X., Sudmant, P.H., Antonacci, F., Graves, T.A., Nefedov, M., Rosenfeld, J.A., Sajjadian, S., Malig, M., Kotkiewicz, H., et al. (2012). Evolution of human-specific neural SRGAP2 genes by incomplete segmental duplication. Cell 149, 912–922. 10.1016/j.cell.2012.03.033

26. Deschamps, C., Faideau, M., Jaber, M., Gaillard, A., and Prestoz, L. (2009). Expression of ephrinA5 during development and potential involvement in the guidance of the mesostriatal pathway. Exp. Neurol. 219, 466–480. 10.1016/j.expneurol.2009.06.020

27. Dixon, J.R., Jung, I., Selvaraj, S., Shen, Y., Antosiewicz-Bourget, J.E., Lee, A.Y., Ye, Z., Kim, A., Rajagopal, N., Xie, W., et al. (2015). Chromatin architecture reorganization during stem cell differentiation. Nature 518, 331–336. 10.1038/nature14222

28. Dodman, N.H., Brown, D.C., and Serpell, J.A. (2018). Associations between owner personality and psychological status and the prevalence of canine behavior problems. PLoS One 13, e0192846. 10.1371/journal.pone.0192846

29. Dodman, N.H., Ginns, E.I., Shuster, L., Moon-Fanelli, A.A., Galdzicka, M., Zheng, J., Ruhe, A.L., and Neff, M.W. (2016). Genomic risk for severe canine compulsive disorder, a dog model of human OCD. Int. J. of Appl. Res. 14.

30. Ducuing, H., Gardette, T., Pignata, A., Tauszig-Delamasure, S., and Castellani, V. (2019). Commissural axon navigation in the spinal cord: A repertoire of repulsive forces is in command. Semin. Cell Dev. Biol. 85, 3–12. 10.1016/j.semcdb.2017.12.010

31. Duffy, D.L., Hsu, Y., and Serpell, J.A. (2008). Breed differences in canine aggression. Appl. Anim. Behav. Sci. 114, 441–460. 10.1016/j.applanim.2008.04.006

32. Duffy, D.L., and Serpell, J.A. (2012). Predictive validity of a method for evaluating temperament in young guide and service dogs. Appl. Anim. Behav. Sci. 138, 99–109. 10.1016/j.applanim.2012.02.011

33. Duncan, J., and Owen, A.M. (2000). Common regions of the human frontal lobe recruited by diverse cognitive demands. Trends Neurosci. 23, 475–483. 10.1016/s0166-2236(00)01633-7

34. Edwards, R.J., Field, M.A., Ferguson, J.M., Dudchenko, O., Keilwagen, J., Rosen, B.D., Johnson, G.S., Rice, E.S., Hillier, L.D., Hammond, J.M., et al. (2021). Chromosome-length genome assembly and structural variations of the primal Basenji dog (Canis lupus familiaris) genome. BMC Genomics 22, 188. 10.1186/s12864-021-07493-6

35. Felsenstein, J. (2005). PHYLIP (Phylogeny Inference Package), version 3.6 (Joseph Felsenstein.).

36. Field, M.A., Rosen, B.D., Dudchenko, O., Chan, E.K.F., Minoche, A.E., Edwards, R.J., Barton, K., Lyons, R.J., Tuipulotu, D.E., Hayes, V.M., et al. (2020). Canfam_GSD: De novo chromosome-length genome assembly of the German Shepherd Dog (Canis lupus familiaris) using a combination of long reads, optical mapping, and Hi-C. GigaScience 9, giaa027. 10.1093/gigascience/giaa027

37. Fleck, M.S., Daselaar, S.M., Dobbins, I.G., and Cabeza, R. (2006). Role of prefrontal and anterior cingulate regions in decision-making processes shared by memory and nonmemory tasks. Cereb. Cortex 16, 1623–1630. 10.1093/cercor/bhj097

38. Fogle, B. (1995). The encyclopedia of the dog (Dorling Kindersley New York).

39. Frantz, L.A., Mullin, V.E., Pionnier-Capitan, M., Lebrasseur, O., Ollivier, M., Perri, A., Linderholm, A., Mattiangeli, V., Teasdale, M.D., Dimopoulos, E.A., et al. (2016). Genomic and archaeological evidence suggest a dual origin of domestic dogs. Science 352, 1228–1231. 10.1126/science.aaf3161

40. Haghverdi, L., Büttner, M., Wolf, F.A., Buettner, F., and Theis, F.J. (2016). Diffusion pseudotime robustly reconstructs lineage branching. Nat. Methods 13, 845–848. 10.1038/nmeth.3971

41. Halo, J.V., Pendleton, A.L., Shen, F., Doucet, A.J., Derrien, T., Hitte, C., Kirby, L.E., Myers, B., Sliwerska, E., Emery, S., et al. (2021). Long-read assembly of a Great Dane genome highlights the contribution of GC-rich sequence and mobile elements to canine genomes. Proc. Natl. Acad. Sci. USA 118, e2016274118. 10.1073/pnas.2016274118

42. Haygood, R., Babbitt, C.C., Fedrigo, O., and Wray, G.A. (2010). Contrasts between adaptive coding and noncoding changes during human evolution. Proc. Natl. Acad. Sci. USA 107, 7853–7857. 10.1073/pnas.0911249107

43. Hayward, J.J., Castelhano, M.G., Oliveira, K.C., Corey, E., Balkman, C., Baxter, T.L., Casal, M.L., Center, S.A., Fang, M., Garrison, S.J., et al. (2016). Complex disease and phenotype mapping in the domestic dog. Nat. Commun. 7, 10460. 10.1038/ncomms10460

44. Hecht, E.E., Smaers, J.B., Dunn, W.D., Kent, M., Preuss, T.M., and Gutman, D.A. (2019). Significant neuroanatomical variation among domestic dog breeds. J. Neurosci. 39, 7748. 10.1523/JNEUROSCI.0303-19.2019

45. Heyne, H.O., Lautenschläger, S., Nelson, R., Besnier, F., Rotival, M., Cagan, A., Kozhemyakina, R., Plyusnina, I.Z., Trut, L., Carlborg, Ö., et al. (2014). Genetic influences on brain gene expression in rats selected for tameness and aggression. Genetics 198, 1277–1290. 10.1534/genetics.114.168948

46. Hoeppner, M.P., Lundquist, A., Pirun, M., Meadows, J.R., Zamani, N., Johnson, J., Sundström, G., Cook, A., FitzGerald, M.G., Swofford, R., et al. (2014). An improved canine genome and a comprehensive catalogue of coding genes and non-coding transcripts. PLoS One 9, e91172. 10.1371/journal.pone.0091172

47. Hsu, Y., and Serpell, J.A. (2003). Development and validation of a questionnaire for measuring behavior and temperament traits in pet dogs. J. Am. Vet. Med. Assoc. 223, 1293–1300. 10.2460/javma.2003.223.1293

48. Ilska, J., Haskell, M.J., Blott, S.C., Sánchez-Molano, E., Polgar, Z., Lofgren, S.E., Clements, D.N., and Wiener, P. (2017). Genetic characterization of dog personality traits. Genetics 206, 1101–1111. 10.1534/genetics.116.192674

49. Jagannathan, V., Drögemüller, C., Leeb, T., and Dog Biomedical Variant Database, C. (2019). A comprehensive biomedical variant catalogue based on whole genome sequences of 582 dogs and eight wolves. Anim. Genet. 50, 695–704. 10.1111/age.12834

50. Jagannathan, V.A.-O., Hitte, C.A.-O.X., Kidd, J.A.-O., Masterson, P., Murphy, T.A.-O., Emery, S., Davis, B.A.-O.X., Buckley, R.A.-O., Liu, Y.H., Zhang, X.Q., et al. (2021). Dog10K_Boxer_Tasha_1.0: A long-read assembly of the dog reference genome. Genes (Basel). 12, 847. 10.3390/genes12060847.

51. Jakuba, T., Polcová, Z., Fedáková, D., Kottferová, J., Mareková, J., Fejsáková, M., Ondrašovičová, O., and Ondrašovič, M. (2013). Differences in evaluation of a dog’s temperament by individual members of the same household. Soc. Anim. 21, 582–589.

52. Jhang, J., Lee, H., Kang, M.S., Lee, H.-S., Park, H., and Han, J.-H. (2018). Anterior cingulate cortex and its input to the basolateral amygdala control innate fear response. Nat. Commun. 9, 2744–2744. 10.1038/s41467-018-05090-y

53. Kang, H.J., Kawasawa, Y.I., Cheng, F., Zhu, Y., Xu, X., Li, M., Sousa, A.M.M., Pletikos, M., Meyer, K.A., Sedmak, G., et al. (2011). Spatio-temporal transcriptome of the human brain. Nature 478, 483–489. 10.1038/nature10523

54. Karlsson, E.K., Baranowska, I., Wade, C.M., Salmon Hillbertz, N.H.C., Zody, M.C., Anderson, N., Biagi, T.M., Patterson, N., Pielberg, G.R., Kulbokas, E.J., et al. (2007). Efficient mapping of mendelian traits in dogs through genome-wide association. Nat. Genet. 39, 1321–1328. 10.1038/ng.2007.10

55. Karlsson Linnér, R., Biroli, P., Kong, E., Meddens, S.F.W., Wedow, R., Fontana, M.A., Lebreton, M., Tino, S.P., Abdellaoui, A., Hammerschlag, A.R., et al. (2019). Genome-wide association analyses of risk tolerance and risky behaviors in over 1 million individuals identify hundreds of loci and shared genetic influences. Nat. Genet. 51, 245–257. 10.1038/s41588-018-0309-3

56. Kim, J., Lee, C., Ko, B.J., Yoo, D., Won, S., Phillippy, A., Fedrigo, O., Zhang, G., Howe, K., Wood, J., et al. (2021). False gene and chromosome losses affected by assembly and sequence errors. bioRxiv, DP9I6

57. Kim, J., Williams, F.J., Dreger, D.L., Plassais, J., Davis, B.W., Parker, H.G., and Ostrander, E.A. (2018). Genetic selection of athletic success in sport-hunting dogs. Proc. Natl. Acad. Sci. USA 115, E7212–e7221. 10.1073/pnas.1800455115

58. Kolde, R. (2015). pheatmap: Pretty Heatmaps. R package version 1.0.8.

59. Kuchroo, M., Huang, J., Wong, P., Grenier, J.-C., Shung, D., Tong, A., Lucas, C., Klein, J., Burkhardt, D.B., Gigante, S., et al. (2022). Multiscale PHATE identifies multimodal signatures of COVID-19. Nat. Biotechnol. 10.1038/s41587-021-01186-x

60. Lane, R.D., Reiman, E.M., Bradley, M.M., Lang, P.J., Ahern, G.L., Davidson, R.J., and Schwartz, G.E. (1997). Neuroanatomical correlates of pleasant and unpleasant emotion. Neuropsychologia 35, 1437–1444. 10.1016/s0028-3932(97)00070-5

61. Larson, G., Karlsson, E.K., Perri, A., Webster, M.T., Ho, S.Y.W., Peters, J., Stahl, P.W., Piper, P.J., Lingaas, F., Fredholm, M., et al. (2012). Rethinking dog domestication by integrating genetics, archeology, and biogeography. Proc. Natl. Acad. Sci. USA 109, 8878–8883. 10.1073/pnas.1203005109

62. Larsson, M. (2015). Binocular vision, the optic chiasm, and their associations with vertebrate motor behavior. Front. Ecol. Evol. 3.

63. Leinonen, R., Sugawara, H., Shumway, M., and International Nucleotide Sequence Database, C. (2011). The sequence read archive. Nucleic acids res. 39, D19–D21. 10.1093/nar/gkq1019

64. Li, H. (2011). A statistical framework for SNP calling, mutation discovery, association mapping and population genetical parameter estimation from sequencing data. Bioinformatics 27, 2987–2993. 10.1093/bioinformatics/btr509

65. Li, H., and Durbin, R. (2009). Fast and accurate short read alignment with Burrows–Wheeler transform. Bioinformatics 25, 1754–1760. 10.1093/bioinformatics/btp324

66. Li, H., Handsaker, B., Wysoker, A., Fennell, T., Ruan, J., Homer, N., Marth, G., Abecasis, G., and Durbin, R. (2009). The Sequence Alignment/Map format and SAMtools. Bioinformatics 25, 2078–2079. 10.1093/bioinformatics/btp352

67. Liberzon, A., Subramanian, A., Pinchback, R., Thorvaldsdóttir, H., Tamayo, P., and Mesirov, J.P. (2011). Molecular signatures database (MSigDB) 3.0. Bioinformatics 27, 1739–1740. 10.1093/bioinformatics/btr260

68. Lindblad-Toh, K., Wade, C.M., Mikkelsen, T.S., Karlsson, E.K., Jaffe, D.B., Kamal, M., Clamp, M., Chang, J.L., Kulbokas, E.J., 3rd, Zody, M.C., et al. (2005). Genome sequence, comparative analysis and haplotype structure of the domestic dog. Nature 438, 803–819. 10.1038/nature04338

69. MacArthur, J., Bowler, E., Cerezo, M., Gil, L., Hall, P., Hastings, E., Junkins, H., McMahon, A., Milano, A., Morales, J., et al. (2017). The new NHGRI-EBI Catalog of published genome-wide association studies (GWAS Catalog). Nucleic acids res. 45, D896–D901. 10.1093/nar/gkw1133

70. MacLean, E.L., Snyder-Mackler, N., vonHoldt, B.M., and Serpell, J.A. (2019). Highly heritable and functionally relevant breed differences in dog behaviour. Proc. Royal Soc. B 286, 20190716. doi:10.1098/rspb.2019.0716

71. Mamiya, P.C., Hennesy, Z., Zhou, R., and Wagner, G.C. (2008). Changes in attack behavior and activity in EphA5 knockout mice. Brain Res. 1205, 91–99. 10.1016/j.brainres.2008.02.047

72. Manichaikul, A., Mychaleckyj, J.C., Rich, S.S., Daly, K., Sale, M., and Chen, W.-M. (2010). Robust relationship inference in genome-wide association studies. Bioinformatics 26, 2867–2873. 10.1093/bioinformatics/btq559

73. McInnes, L., Healy, J., Saul, N., and Großberger, L. (2018). Umap: uniform manifold approximation and projection, J. Open Source Softw. 3, 861.

74. McLaren, W., Gil, L., Hunt, S.E., Riat, H.S., Ritchie, G.R.S., Thormann, A., Flicek, P., and Cunningham, F. (2016). The Ensembl Variant Effect Predictor. Genome Biology 17, 122. 10.1186/s13059-016-0974-4

75. McLean, C.Y., Bristor, D., Hiller, M., Clarke, S.L., Schaar, B.T., Lowe, C.B., Wenger, A.M., and Bejerano, G. (2010). GREAT improves functional interpretation of cis-regulatory regions. Nat. Biotechnol. 28, 495–501. 10.1038/nbt.1630

76. Mech, L.D., Smith, D.W., and MacNulty, D.R. (2015). Wolves on the Hunt (University of Chicago Press).

77. Moon, K.R., van Dijk, D., Wang, Z., Gigante, S., Burkhardt, D.B., Chen, W.S., Yim, K., Elzen, A.v.d., Hirn, M.J., Coifman, R.R., et al. (2019). Visualizing structure and transitions in high-dimensional biological data. Nat. Biotechnol. 37, 1482–1492. 10.1038/s41587-019-0336-3

78. Moore, J.E., Purcaro, M.J., Pratt, H.E., Epstein, C.B., Shoresh, N., Adrian, J., Kawli, T., Davis, C.A., Dobin, A., Kaul, R., et al. (2020). Expanded encyclopaedias of DNA elements in the human and mouse genomes. Nature 583, 699–710. 10.1038/s41586-020-2493-4

79. Morrill, K., Hekman, J., Li, X., McClure, J., Logan, B., Goodman, L., Gao, M., Dong, Y., Alonso, M., Carmichael, E., et al. (2022). Ancestry-inclusive dog genomics challenges popular breed stereotypes. Science 376. 10.1126/science.abk0639

80. Nagasawa, M., Tsujimura, A., Tateishi, K., Mogi, K., Ohta, M., Serpell, J.A., and Kikusui, T. (2011). Assessment of the factorial structures of the C-BARQ in Japan. J. Vet. Med. Sci. 73, 869–875. 10.1292/jvms.10-0208

81. Naval-Sanchez, M., Nguyen, Q., McWilliam, S., Porto-Neto, L.R., Tellam, R., Vuocolo, T., Reverter, A., Perez-Enciso, M., Brauning, R., Clarke, S., et al. (2018). Sheep genome functional annotation reveals proximal regulatory elements contributed to the evolution of modern breeds. Nat. Commun. 9, 859. 10.1038/s41467-017-02809-1

82. Niepoth, N., and Bendesky, A. (2020). How natural genetic variation shapes behavior. Ann. Rev. Genomics and Hum. Genet. 21, 437–463. 10.1146/annurev-genom-111219-080427

83. Ostrander, E.A., Wang, G.-D., Larson, G., vonHoldt, B.M., Davis, B.W., Jagannathan, V., Hitte, C., Wayne, R.K., Zhang, Y.-P., and Dog, K.C. (2019). Dog10K: an international sequencing effort to advance studies of canine domestication, phenotypes and health. Natl. Sci. Rev. 6, 810–824. 10.1093/nsr/nwz049

84. Parker, H.G., Dreger, D.L., Rimbault, M., Davis, B.W., Mullen, A.B., Carpintero-Ramirez, G., and Ostrander, E.A. (2017). Genomic analyses reveal the influence of geographic origin, migration, and hybridization on modern dog breed development. Cell Rep. 19, 697–708. 10.1016/j.celrep.2017.03.079

85. Parker, H.G., Kim, L.V., Sutter, N.B., Carlson, S., Lorentzen, T.D., Malek, T.B., Johnson, G.S., DeFrance, H.B., Ostrander, E.A., and Kruglyak, L. (2004). Genetic structure of the purebred domestic dog. Science 304, 1160–1164. 10.1126/science.1097406

86. Pendleton, A.L., Shen, F., Taravella, A.M., Emery, S., Veeramah, K.R., Boyko, A.R., and Kidd, J.M. (2018). Comparison of village dog and wolf genomes highlights the role of the neural crest in dog domestication. BMC Biol. 16, 64. 10.1186/s12915-018-0535-2

87. Perri, A.R., Feuerborn, T.R., Frantz, L.A.F., Larson, G., Malhi, R.S., Meltzer, D.J., and Witt, K.E. (2021). Dog domestication and the dual dispersal of people and dogs into the Americas. Proc. Natl. Acad. Sci. USA 118, e2010083118. 10.1073/pnas.2010083118

88. Plassais, J., Kim, J., Davis, B.W., Karyadi, D.M., Hogan, A.N., Harris, A.C., Decker, B., Parker, H.G., and Ostrander, E.A. (2019). Whole genome sequencing of canids reveals genomic regions under selection and variants influencing morphology. Nat. Commun. 10, 1489. 10.1038/s41467-019-09373-w

89. Plassais, J., Rimbault, M., Williams, F.J., Davis, B.W., Schoenebeck, J.J., and Ostrander, E.A. (2017). Analysis of large versus small dogs reveals three genes on the canine X chromosome associated with body weight, muscling and back fat thickness. PLoS Genet. 13, e1006661. 10.1371/journal.pgen.1006661

90. Pollard, K.S., Hubisz, M.J., Rosenbloom, K.R., and Siepel, A. (2010). Detection of nonneutral substitution rates on mammalian phylogenies. Genome Res. 20, 110–121. 10.1101/gr.097857.109

91. Prieur, D.S., and Rebsam, A. (2017). Retinal axon guidance at the midline: Chiasmatic misrouting and consequences. Dev. Neurobiol. 77, 844–860.

92. Purcell, S., Neale, B., Todd-Brown, K., Thomas, L., Ferreira, M.A.R., Bender, D., Maller, J., Sklar, P., de Bakker, P.I.W., Daly, M.J., et al. (2007). PLINK: a tool set for whole-genome association and population-based linkage analyses. Am. J. Hum. Genet. 81, 559–575. 10.1086/519795

93. Rhead, B., Karolchik, D., Kuhn, R.M., Hinrichs, A.S., Zweig, A.S., Fujita, P.A., Diekhans, M., Smith, K.E., Rosenbloom, K.R., Raney, B.J., et al. (2010). The UCSC Genome Browser database: update 2010. Nucleic Acids Res. 38, D613–619. 10.1093/nar/gkp939

94. Rhie, A., McCarthy, S.A., Fedrigo, O., Damas, J., Formenti, G., Koren, S., Uliano-Silva, M., Chow, W., Fungtammasan, A., Kim, J., et al. (2021). Towards complete and error-free genome assemblies of all vertebrate species. Nature 592, 737–746. 10.1038/s41586-021-03451-0

95. Rimbault, M., Beale, H.C., Schoenebeck, J.J., Hoopes, B.C., Allen, J.J., Kilroy-Glynn, P., Wayne, R.K., Sutter, N.B., and Ostrander, E.A. (2013). Derived variants at six genes explain nearly half of size reduction in dog breeds. Genome Res. 23, 1985–1995. 10.1101/gr.157339.113

96. Sabeti, P.C., Schaffner, S.F., Fry, B., Lohmueller, J., Varilly, P., Shamovsky, O., Palma, A., Mikkelsen, T.S., Altshuler, D., and Lander, E.S. (2006). Positive natural selection in the human lineage. Science 312, 1614–1620. 10.1126/science.1124309

97. Salonen, M., Sulkama, S., Mikkola, S., Puurunen, J., Hakanen, E., Tiira, K., Araujo, C., and Lohi, H. (2020). Prevalence, comorbidity, and breed differences in canine anxiety in 13,700 Finnish pet dogs. Sci. Rep. 10, 1–11.

98. Sánchez-Villagra, M.R., Geiger, M., and Schneider, R.A. The taming of the neural crest: a developmental perspective on the origins of morphological covariation in domesticated mammals. R. Soc. Open Sci. 3, 160107. 10.1098/rsos.160107

99. Savolainen, P., Zhang, Y.P., Luo, J., Lundeberg, J., and Leitner, T. (2002). Genetic evidence for an East Asian origin of domestic dogs. Science 298, 1610–1613. 10.1126/science.1073906

100. Schellinck, H.M., Cyr, D.P., Brown, R.E. (2010). Chapter 7 - How many ways can mouse behavioral experiments go wrong? Confounding variables in mouse models of neurodegenerative diseases and how to control them. In Advances in the Study of Behavior, Vol 41, H.J. Brockmann, T.J. Roper, M. Naguib, K.E. Wynne-Edwards, J.C. Mitani, L.W. Simmonsm, eds. (Academic Press), pp. 255–366.

101. Schlamp, F., van der Made, J., Stambler, R., Chesebrough, L., Boyko, A.R., and Messer, P.W. (2016). Evaluating the performance of selection scans to detect selective sweeps in domestic dogs. Mol. Ecol. 25, 342–356. 10.1111/mec.13485

102. Schoenebeck, J.J., Hamilton-Dyer, S., Baxter, I.L., Schwarz, T., and Nussbaumer, M. (2021). From head to hind: Elucidating function through contrasting morphometrics of ancient and modern pedigree dogs. Anat. Rec. (Hoboken) 304, 63–77. 10.1002/ar.24412

103. Scott, J.P., and Fuller, J.L. (2012). Genetics and the Social Behavior of the Dog, Vol 570 (University of Chicago Press).

104. Serpell, J. (2017). The Domestic Dog (Cambridge University Press).

105. Serpell, J.A., and Duffy, D.L. (2014). Dog breeds and their behavior. In Domestic Dog Cognition and Behavior: The Scientific Study of Canis familiaris, A. Horowitz, ed. (Berlin, Heidelberg: Springer Berlin Heidelberg), pp. 31–57.

106. Serpell, J.A., and Hsu, Y.A. (2005). Effects of breed, sex, and neuter status on trainability in dogs. Anthrozoös 18, 196–207. 10.2752/089279305785594135

107. Serres-Armero, A., Davis, B.W., Povolotskaya, I.S., Morcillo-Suarez, C., Plassais, J., Juan, D., Ostrander, E.A., and Marques-Bonet, T. (2021). Copy number variation underlies complex phenotypes in domestic dog breeds and other canids. Genome res. 31, 762–774. 10.1101/gr.266049.120

108. Sheleg, M., Yu, Q., Go, C., Wagner, G.C., Kusnecov, A.W., and Zhou, R. (2017). Decreased maternal behavior and anxiety in ephrin-A5(-/-) mice. Genes Brain Behav. 16, 271–284. 10.1111/gbb.12319

109. Sinding, M.-H.S., Gopalakrishnan, S., Ramos-Madrigal, J., de Manuel, M., Pitulko, V.V., Kuderna, L., Feuerborn, T.R., Frantz, L.A.F., Vieira, F.G., Niemann, J., et al. (2020). Arctic-adapted dogs emerged at the Pleistocene–Holocene transition. Science 368, 1495–1499. 10.1126/science.aaz8599

110. Snow, P.J. (2016). The structural and functional organization of cognition. Front. Hum. Neurosci. 10, 501–501. 10.3389/fnhum.2016.00501

111. Street, K., Risso, D., Fletcher, R.B., Das, D., Ngai, J., Yosef, N., Purdom, E., and Dudoit, S. (2018). Slingshot: cell lineage and pseudotime inference for single-cell transcriptomics. BMC Genomics 19, 477. 10.1186/s12864-018-4772-0

112. Sutter, N.B., Bustamante, C.D., Chase, K., Gray, M.M., Zhao, K., Zhu, L., Padhukasahasram, B., Karlins, E., Davis, S., Jones, P.G., et al. (2007). A Single IGF1 allele is a major determinant of small size in dogs. Science 316, 112–115. 10.1126/science.1137045

113. Svartberg, K. (2006). Breed-typical behaviour in dogs—Historical remnants or recent constructs? Appl. Anim. Behav. Sci. 96, 293–313. 10.1016/j.applanim.2005.06.014

114. Talenti, A., Dreger, D.L., Frattini, S., Polli, M., Marelli, S., Harris, A.C., Liotta, L., Cocco, R., Hogan, A.N., Bigi, D., et al. (2018). Studies of modern Italian dog populations reveal multiple patterns for domestic breed evolution. Ecol. Evol. 8, 2911–2925. 10.1002/ece3.3842

115. Tang, R., Noh, H.J., Wang, D., Sigurdsson, S., Swofford, R., Perloski, M., Duxbury, M., Patterson, E.E., Albright, J., Castelhano, M., et al. (2014). Candidate genes and functional noncoding variants identified in a canine model of obsessive-compulsive disorder. Genome Biol. 15, R25. 10.1186/gb-2014-15-3-r25

116. Thalmann, O., Shapiro, B., Cui, P., Schuenemann, V.J., Sawyer, S.K., Greenfield, D.L., Germonpré, M.B., Sablin, M.V., López-Giráldez, F., Domingo-Roura, X., et al. (2013). Complete mitochondrial genomes of ancient canids suggest a European origin of domestic dogs. Science 342, 871–874. 10.1126/science.1243650

117. Turner, K.M., and Parkes, S.L. (2020). Prefrontal regulation of behavioural control: Evidence from learning theory and translational approaches in rodents. Neurosci. Biobehav. Rev. 118, 27–41. 10.1016/j.neubiorev.2020.07.010

118. Van der Auwera, G.A., Carneiro, M.O., Hartl, C., Poplin, R., Del Angel, G., Levy-Moonshine, A., Jordan, T., Shakir, K., Roazen, D., Thibault, J., et al. (2013). From FastQ data to high confidence variant calls: the Genome Analysis Toolkit best practices pipeline. Curr. Protoc. Bioinformatics 43, 11.10.11–11.10.33. 10.1002/0471250953.bi1110s43

119. van Rooy, D., Arnott, E.R., Early, J.B., McGreevy, P., and Wade, C.M. (2014). Holding back the genes: limitations of research into canine behavioural genetics. Canine Genet. Epidemiol. 1, 7–7. 10.1186/2052-6687-1-7

120. Vasung, L., Huang, H., Jovanov-Milošević, N., Pletikos, M., Mori, S., and Kostović, I. (2010). Development of axonal pathways in the human fetal fronto-limbic brain: histochemical characterization and diffusion tensor imaging. J. Anat. 217, 400–417. 10.1111/j.1469-7580.2010.01260.x

121. Vaysse, A., Ratnakumar, A., Derrien, T., Axelsson, E., Rosengren Pielberg, G., Sigurdsson, S., Fall, T., Seppälä, E.H., Hansen, M.S.T., Lawley, C.T., et al. (2011). Identification of genomic regions associated with phenotypic variation between dog breeds using selection mapping. PLOS Genetics 7, e1002316. 10.1371/journal.pgen.1002316

122. vonHoldt, B.M., Pollinger, J.P., Lohmueller, K.E., Han, E., Parker, H.G., Quignon, P., Degenhardt, J.D., Boyko, A.R., Earl, D.A., Auton, A., et al. (2010). Genome-wide SNP and haplotype analyses reveal a rich history underlying dog domestication. Nature 464, 898–902. 10.1038/nature08837

123. vonHoldt, B.M., Shuldiner, E., Koch Ilana, J., Kartzinel Rebecca, Y., Hogan, A., Brubaker, L., Wanser, S., Stahler, D., Wynne Clive, D.L., Ostrander Elaine, A., et al. (2017). Structural variants in genes associated with human Williams-Beuren syndrome underlie stereotypical hypersociability in domestic dogs. Sci. Adv. 3, e1700398. 10.1126/sciadv.1700398

124. Wang, C., Wallerman, O., Arendt, M.-L., Sundström, E., Karlsson, Å., Nordin, J., Mäkeläinen, S., Pielberg, G.R., Hanson, J., Ohlsson, Å., et al. (2021). A novel canine reference genome resolves genomic architecture and uncovers transcript complexity. Commun. Biol. 4, 185. 10.1038/s42003-021-01698-x

125. Wang, X., Pipes, L., Trut, L.N., Herbeck, Y., Vladimirova, A.V., Gulevich, R.G., Kharlamova, A.V., Johnson, J.L., Acland, G.M., Kukekova, A.V., et al. (2018). Genomic responses to selection for tame/aggressive behaviors in the silver fox (Vulpes vulpes). Proc. Natl. Acad. Sci. USA 115, 10398–10403. 10.1073/pnas.1800889115

126. Watanabe, K., Taskesen, E., van Bochoven, A., and Posthuma, D. (2017). Functional mapping and annotation of genetic associations with FUMA. Nat. Commun. 8, 1826

127. Wayne, R.K., and vonHoldt, B.M. (2012). Evolutionary genomics of dog domestication. Mamm. Genome 23, 3–18. 10.1007/s00335-011-9386-7

128. Weber, E.M., and Olsson, I.A.S. (2008). Maternal behaviour in Mus musculus sp.: An ethological review. Appl. Anim. Behav. Sci. 114, 1–22. 10.1016/j.applanim.2008.06.006

129. Webster, M.T., Kamgari, N., Perloski, M., Hoeppner, M.P., Axelsson, E., Hedhammar, Å., Pielberg, G., and Lindblad-Toh, K. (2015). Linked genetic variants on chromosome 10 control ear morphology and body mass among dog breeds. BMC Genomics 16, 474. 10.1186/s12864-015-1702-2

130. Wells, D.L., Morrison, D.J., and Hepper, P.G. (2012). The effect of priming on perceptions of dog breed traits. Anthrozoös 25, 369–377. 10.2752/175303712X13403555186370

131. Wickham, H. (2016). ggplot2: Elegant Graphics for Data Analysis.

132. Wilcox, B., and Walkowicz, C. (1989). Atlas of Dog Breeds of the World. New rev.

133. Wilkins, A.S., Wrangham, R.W., and Fitch, W.T. (2014). The “Domestication Syndrome” in mammals: A unified explanation based on neural crest cell behavior and genetics. Genetics 197, 795–808. 10.1534/genetics.114.165423

134. Willsey, A.J., Sanders, Stephan J., Li, M., Dong, S., Tebbenkamp, Andrew T., Muhle, Rebecca A., Reilly, Steven K., Lin, L., Fertuzinhos, S., Miller, Jeremy A., et al. (2013). Coexpression networks implicate human midfetal deep cortical projection neurons in the pathogenesis of autism. Cell 155, 997–1007. 10.1016/j.cell.2013.10.020

135. Won, H., Huang, J., Opland, C.K., Hartl, C.L., and Geschwind, D.H. (2019). Human evolved regulatory elements modulate genes involved in cortical expansion and neurodevelopmental disease susceptibility. Nat. Commun. 10, 2396. 10.1038/s41467-019-10248-3

136. Worboys, M., Strange, J.-M., and Pemberton, N. (2018). The Invention of the Modern Dog: Breed and Blood in Victorian Britain (Johns Hopkins University Press).

137. Zapata, I., Serpell, J.A., and Alvarez, C.E. (2016). Genetic mapping of canine fear and aggression. BMC Genomics 17, 572. 10.1186/s12864-016-2936-3

138. Zarrei, M., Burton, C.L., Engchuan, W., Young, E.J., Higginbotham, E.J., MacDonald, J.R., Trost, B., Chan, A.J.S., Walker, S., Lamoureux, S., et al. (2019). A large data resource of genomic copy number variation across neurodevelopmental disorders. npj Genom. Med. 4. 10.1038/s41525-019-0098-3

139. Zhou, J., and Troyanskaya, O.G. (2015). Predicting effects of noncoding variants with deep learning–based sequence model. Nat. Methods 12, 931–934. 10.1038/nmeth.3547

