## Supplementary figures and images for "Domestic dog lineages reveal genetic drivers of behavioral diversification"

### Supplemental Figure 1

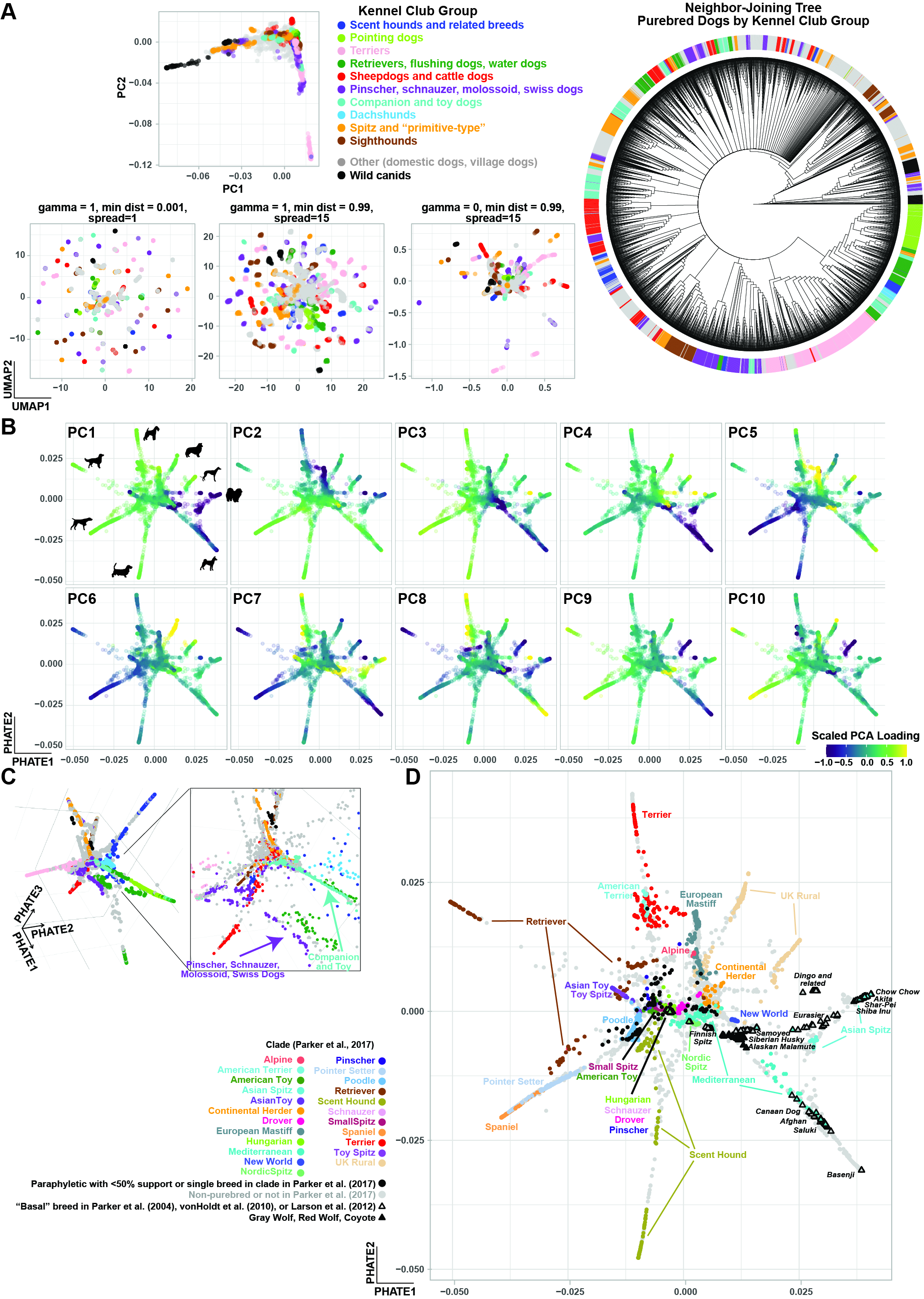

### Supplemental Figure 2

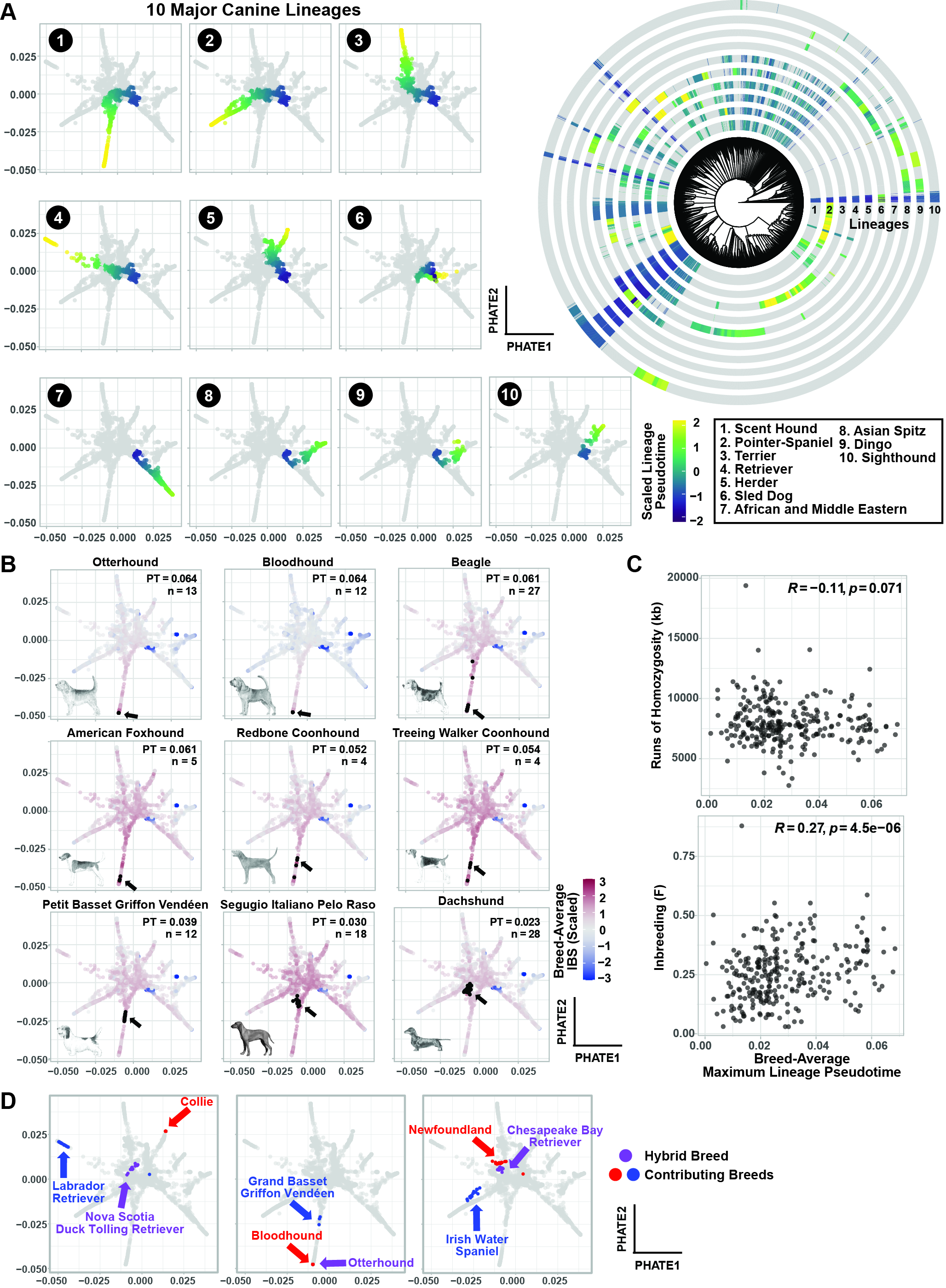

### Supplemental Figure 3

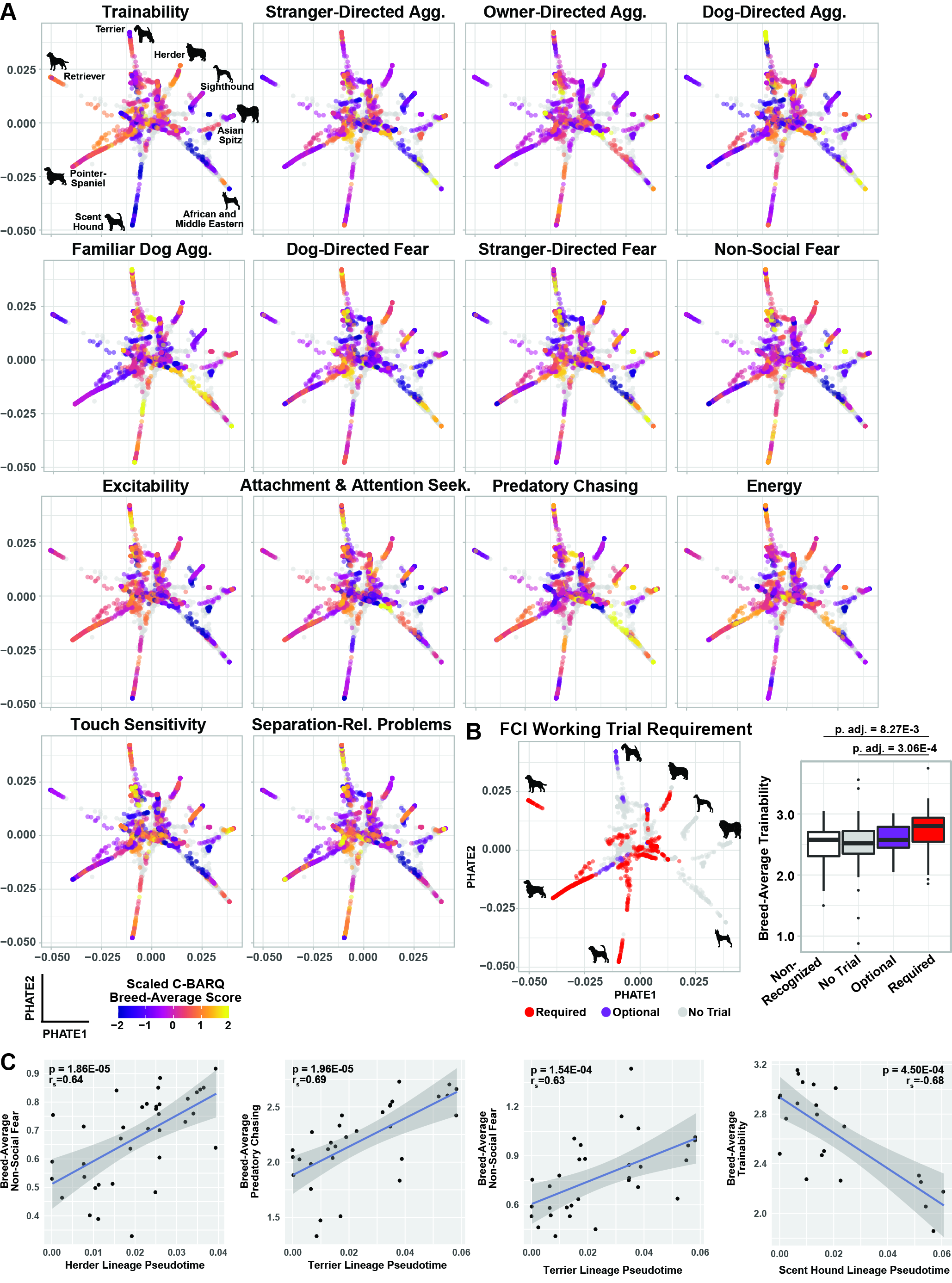

### Supplemental Figure 4

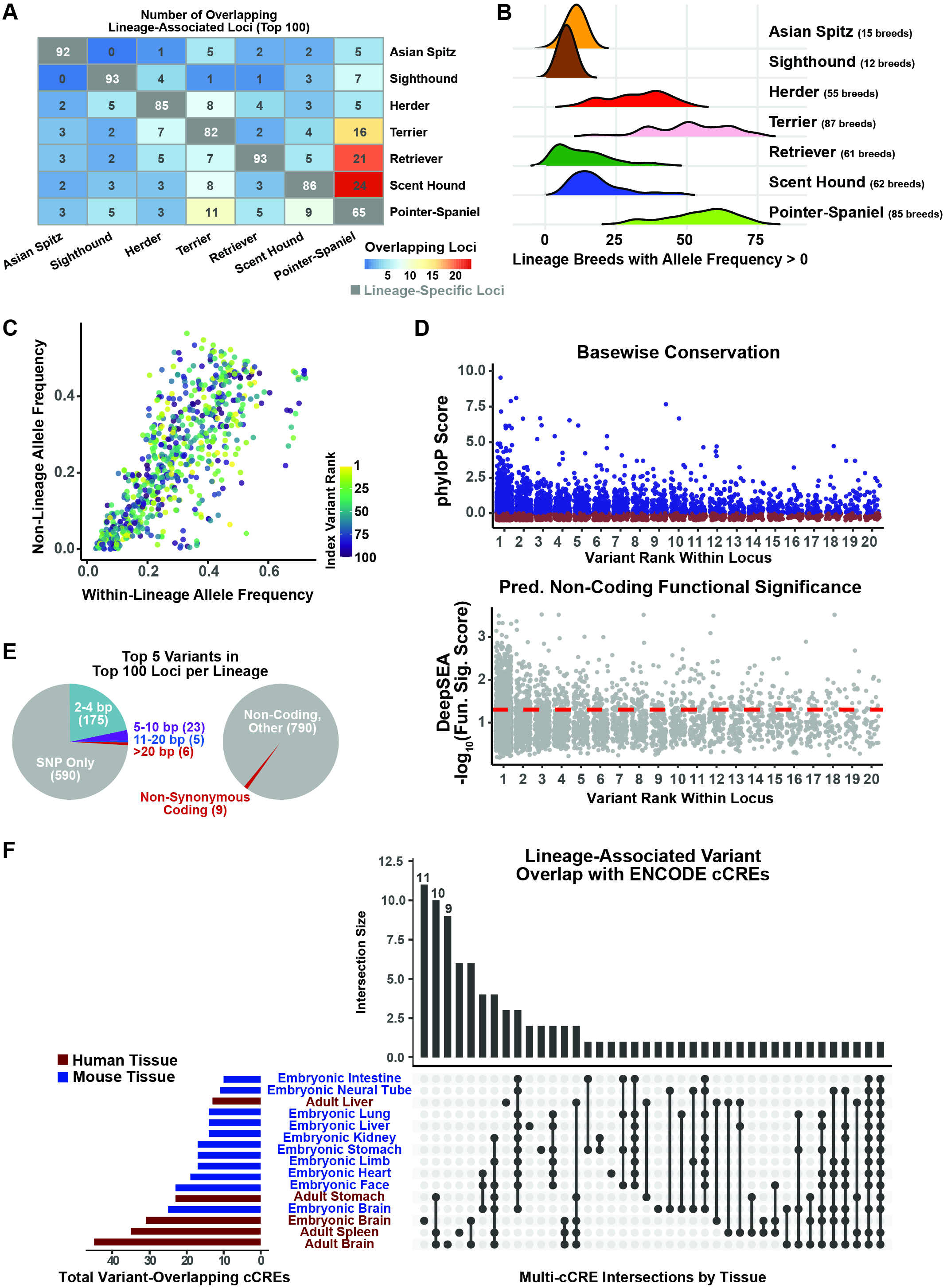

### Supplemental Figure 5

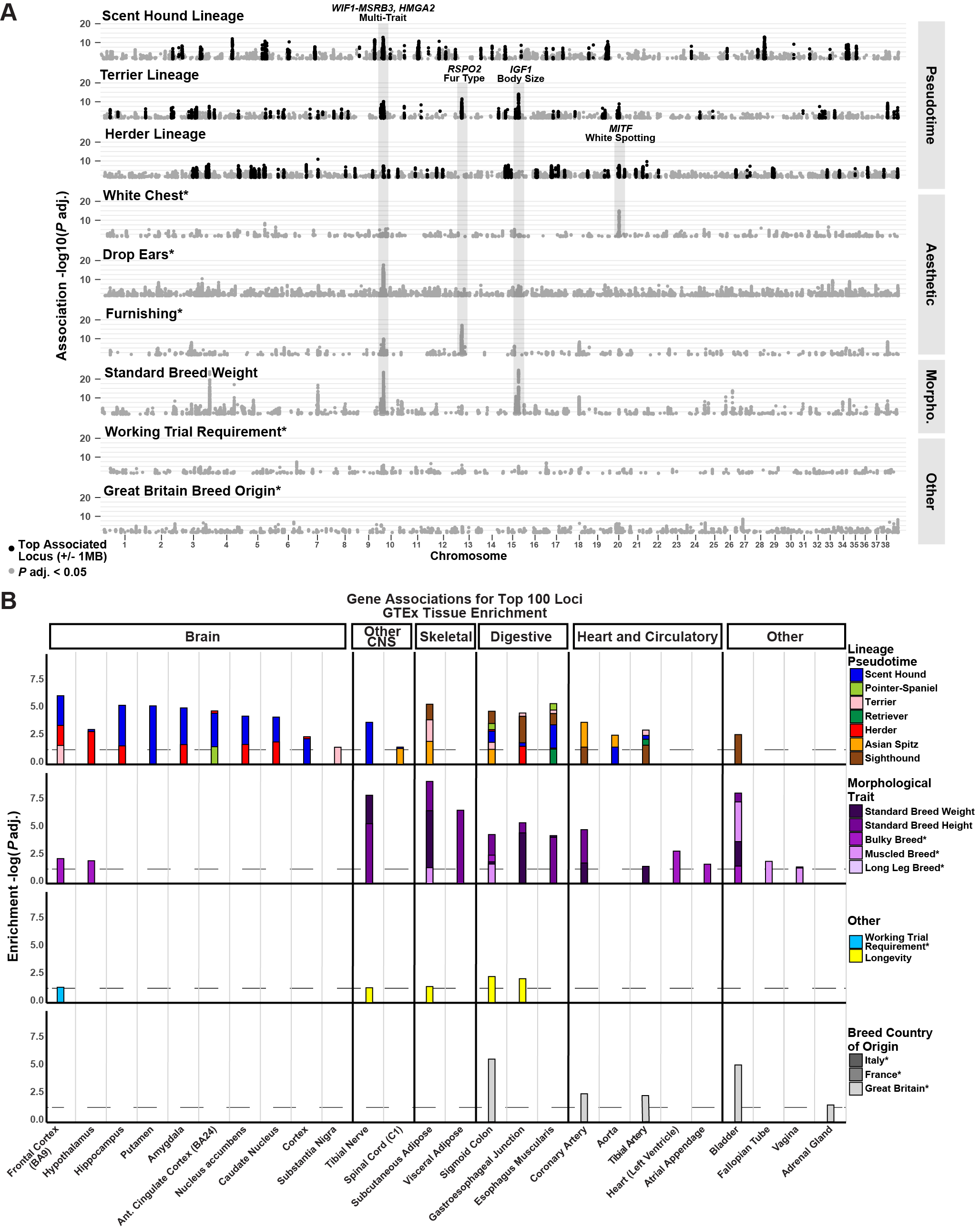

### Supplemental Figure 6

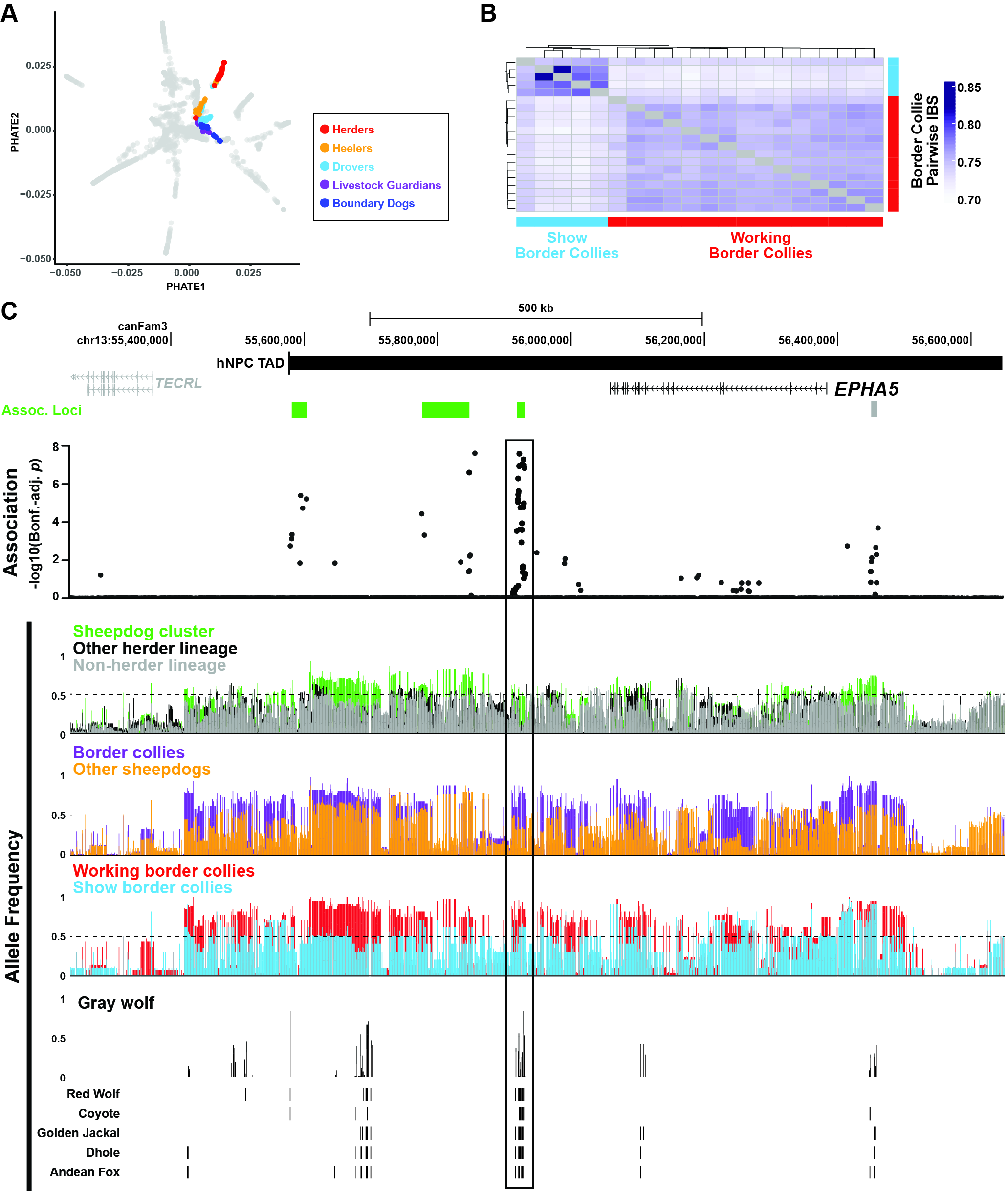
